# Trait misalignment risk in North American forests under climate change

**DOI:** 10.64898/2026.03.13.711509

**Authors:** Alastair Pickering, Tim Newbold, Alex L. Pigot, Carolina Tovar, Daniel S. Maynard

## Abstract

Climate change is expected to alter forest community composition and functioning, with consequences for the ecosystem services forests provide. However, most macroecological projections focus on individual species distributions and offer limited insight into whether entire communities will remain functionally compatible with future climatic conditions. Here we quantify the risk that present-day forest communities will become functionally misaligned with projected climates using a trait-based approach. We analysed forest inventory data from more than 42,000 mature plots across the United States and Canada. For each plot we estimated community-weighted means for 24 functional traits describing leaf economics, hydraulic function, wood structure, abiotic tolerances and symbiotic strategies. We modelled relationships between community functional composition and environmental conditions, and used these relationships to estimate the trait profiles most compatible with projected late-century climates (2080–2100). Trait–environment misalignment (TEM) risk was quantified as the multivariate distance between current community trait composition and the trait profile associated with the projected future climate at each location, accounting for covariance among traits and intraspecific trait variation. Projected climatic conditions favour trait combinations associated with greater hydraulic capacity and reduced cold and shade tolerance. However, the magnitude of functional misalignment varies strongly across space. The highest TEM risk occurs in high-latitude and montane conifer forests across western and central North America, whereas many mid-latitude broadleaf and mixed forests show lower risk because projected climatic changes reinforce existing drought-adapted functional strategies. Critically, high species richness was the strongest predictor of reduced risk, reinforcing the importance of biodiversity in buffering against adverse outcomes. Our results suggest that many forests are projected to experience climatic conditions associated with functional strategies that differ from those characterising the current community. By identifying where the largest functional adjustments are implied, this trait-based framework provides a scalable way to pinpoint forests most likely to experience suboptimal climate conditions and to prioritise monitoring and climate-adapted management.

## Introduction

Anthropogenic climate change is fundamentally reshaping ecological communities, altering their composition, structure, and function, and threatening their long-term stability (Pecl *et al*., 2017). These shifts have profound implications for the estimated $44-$145 trillion in annual economic value tied to ecosystem services (Herweijer *et al*., 2020; FAO, 2022). Forests, which contribute over $7 trillion per annum to global Gross Domestic Product, play a crucial role in delivering these services, including timber production, carbon sequestration, and water regulation (FAO, 2022). Mounting evidence suggests that climate change is increasingly disrupting wooded ecosystems, with more than 30% of tree species currently threatened with extinction (Rivers *et al*., 2023). Because trees are long-lived, many existing individuals and communities will persist through rapid climatic shifts, with consequences for stability, productivity and the services they provide (Morin *et al*., 2018). A key management challenge therefore is to pinpoint where today’s forest communities are at risk of being functionally incompatible with future climates in order to target adaptive management that maintains functional integrity and essential services (Lindner *et al*., 2014; Keenan, 2015).

A common approach to predicting the community-level impact of environmental change is to establish statistical correlations between observed species distributions and environmental conditions (e.g., joint-species distribution models) (Pollock *et al*., 2014; Norberg *et al*., 2019). While effective for large-scale biodiversity projections of individual taxa, these so-called ‘bottom-up’ approaches encounter several challenges when the aim is to assess whether existing communities will remain similar as climates change (Poggiato *et al*., 2021). Most notably, taxonomic approaches predict probabilities of individual species’ occurrences under future conditions, but these remain confined to the species level and offer limited indication of whether a community—viewed as a top-down, coherent ecological unit—will persist in a comparable functional state (Poggiato *et al*., 2021). Communities may experience taxonomic turnover while maintaining cohesion, particularly when lost species are replaced by functionally similar ones (Rosenfeld, 2002). These projections also do not directly indicate whether key community-level functions will be maintained, especially where functional redundancy can sustain processes despite compositional change (Cadotte, Carscadden and Mirotchnick, 2011). Furthermore, scaling species-level predictions to the community level is computationally intensive even at moderate levels of species richness (Wilkinson *et al*., 2021).

An alternate approach to identifying communities at risk of functional change is to focus on top-down, community-level properties—such as biomass, productivity, and functional composition—that scale predictably with climate (Michaletz *et al*., 2014; Enquist *et al*., 2015). In particular, community-level traits summarise functional composition while capturing key mechanisms underpinning persistence, functioning, and resilience (Muscarella and Uriarte, 2016; Miller, Damschen and Ives, 2019; Kandlikar, Kleinhesselink and Kraft, 2022). These community functional traits can be used to detect early shifts in community composition, potentially signalling ecological stress before major species losses occur (Andrew *et al*., 2022). By compressing species-level information into a small set of interpretable dimensions, trait-based approaches also enable computationally efficient analyses of broad-scale variation in functional composition. Critically, trait estimates derived from large databases carry quantifiable uncertainty, which can be propagated through downstream analyses to bound predictions and capture the range of plausible functional variation within and across species (Schrodt *et al*., 2015). Moreover, by reflecting the functional strategies that prevail under a given environment, community-level trait distributions provide a baseline for determining how well the resident community is aligned with future climate.

Trait–environment relationships describe how community-level functional composition varies with climate, soils and topography, and therefore which trait profiles are commonly associated with particular environmental conditions (Michaletz *et al*., 2014; Enquist *et al*., 2015; Muscarella and Uriarte, 2016). As climates change, long-lived forests can become increasingly out of step with the conditions they experience, because their existing trait profiles reflect past environments and may not match the trait profiles typically associated with the new climate (Green *et al*., 2022) (see conceptual Fig. 1). The magnitude of this misalignment reflects the functional adjustment implied for the community to converge on the functional composition generally observed under future conditions. Such adjustment can arise through intraspecific trait variation and phenotypic plasticity within existing species, or through demographic processes such as recruitment, turnover and succession that alter dominance and composition (Lajoie and Vellend, 2018). In long-lived forests, legacy cohorts can slow demographic change, allowing misalignment to persist and extend departures in growth, productivity, or resilience (Lamarque *et al*., 2014; Hevia *et al*., 2017). Larger misalignment therefore implies a lower likelihood that the community will retain its present functional composition, while the extent of taxonomic change depends on the balance between within-species responses and demographic turnover. Identifying where misalignment risk is greatest can therefore pinpoint where the largest adjustment is implied, and where management interventions such as monitoring and experimentation are most warranted.

**Figure 1.**
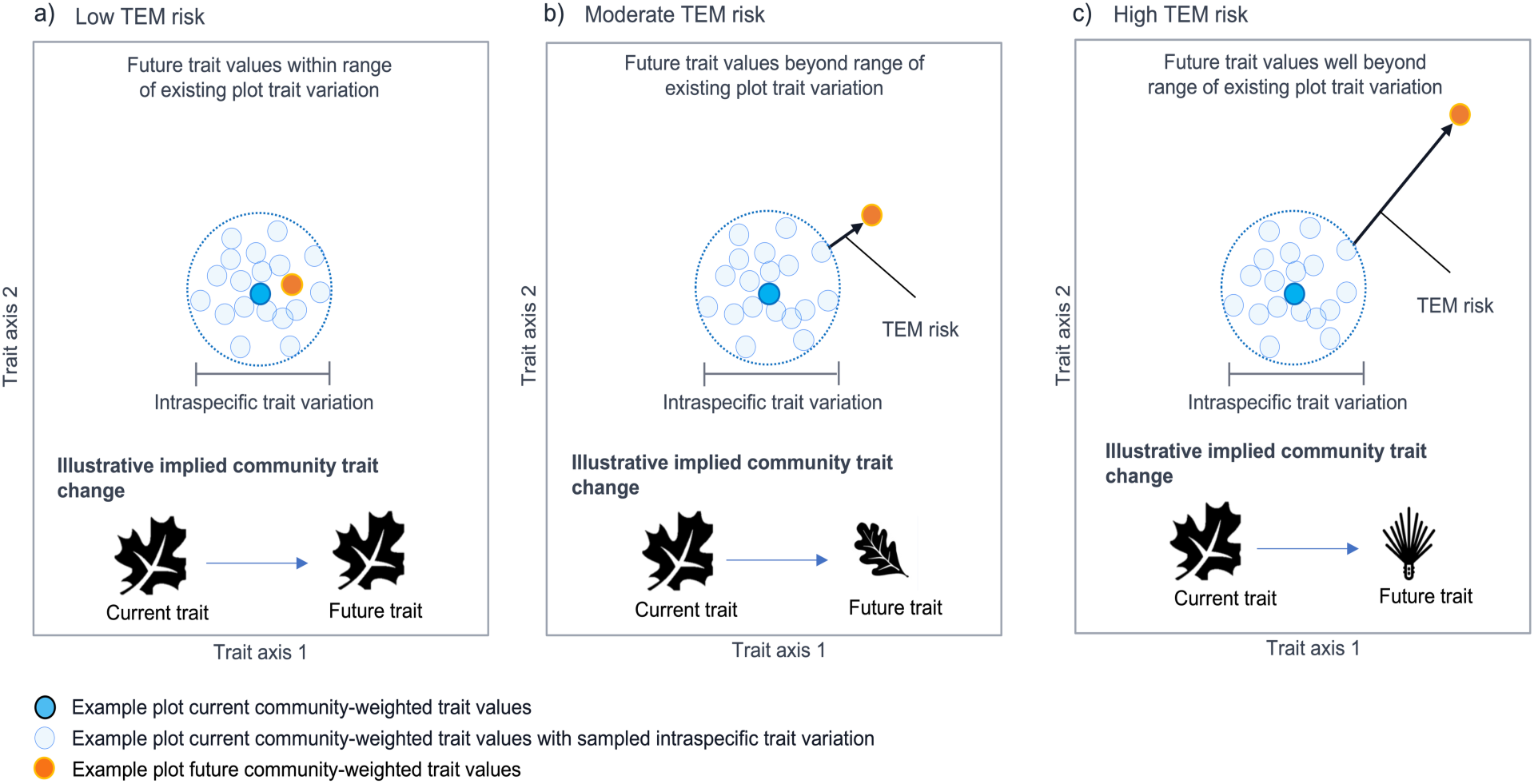
Conceptual illustration of Trait–Environment Misalignment (TEM) risk. Each panel shows the community-weighted trait profile of a single forest plot projected into two-dimensional trait space. The dark blue point represents the present-day community-weighted trait values of the plot, while the pale blue points capture the range of plausible trait values accounting for intraspecific trait variation. The dashed blue circle delineates the 95th percentile of this variation. The orange point represents the predicted future trait values for the same plot, taken as the mean of 500 bootstrap replicates incorporating sampling uncertainty. In panels b and c, the arrow denotes the TEM risk value, defined as the minimum Mahalanobis distance between the modelled future trait profile and the distribution of current trait values (sampled intraspecific variation). In panel a, the future trait values fall within the range of current trait variation and therefore indicate low TEM risk. In panel b, the future values lie outside the current variation but remain close to it in trait space, representing moderate TEM risk. In panel c, the future values lie far beyond the current range of trait variation, indicating a high risk of trait–environment misalignment.

**Figure 2.**
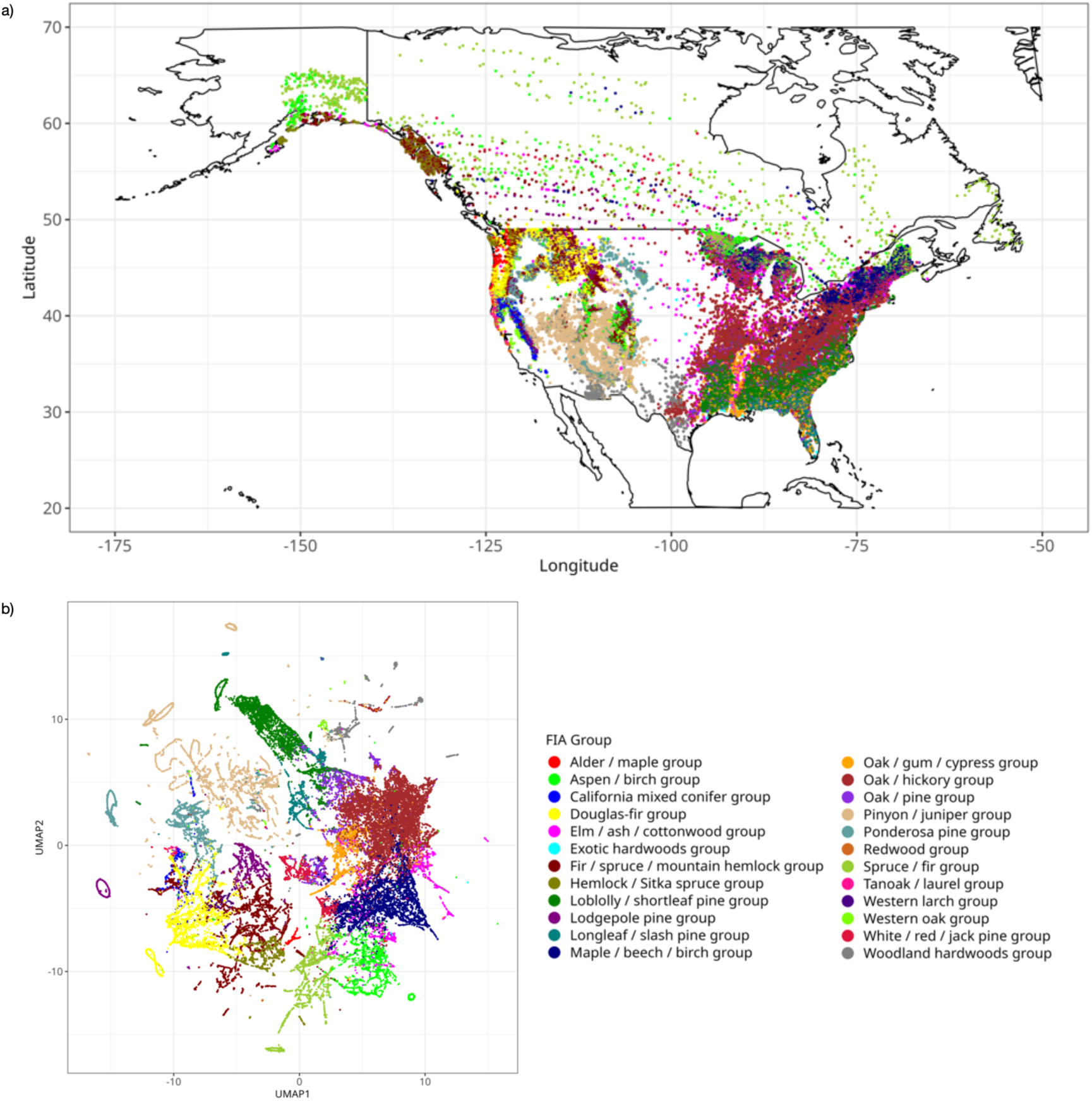
Geographic and trait-space distribution of North American forest plots and their groups. a) Map of the > 42 000 mature forest inventory plots. b) 2D depiction of the 24 plot-level traits across FIA taxonomic groups (using UMAP cosine distance; n neighbors = 5; min_dist = 0.2), illustrating how plots from the same taxonomic group tend to occupy similar regions of trait space. Colours denote FIA group classifications.

Such interventions are especially relevant in heavily managed, commercial ecosystems where direct interventions—like assisted migration and targeted species introductions (Pedlar *et al*., 2012; Palik *et al*., 2022)—are feasible. North American forests, among the most intensively managed and commercially significant wooded ecosystems globally (Campbell and Brown, 2012), span extensive ecological and climatic gradients that provide an ideal backdrop for exploring climate impacts. In these systems, rising temperatures are expected to reduce water availability and intensify drought conditions (see Supplementary Fig. S1 for projected North American environmental changes), factors that may heighten exposure and stress in communities experiencing severe water stress (Mildrexler *et al*., 2016). At the same time, the naturally high species richness in some regions may enhance functional diversity, potentially buffering communities against these climatic pressures (McCann, 2000). Against this backdrop, a key gap is the lack of a scalable framework for identifying where current forest functional composition is likely to become misaligned with projected future climates, hindering efforts to implement climate-adapted forest management.

Here we develop and apply a trait-based metric of trait–environment misalignment (TEM) risk that incorporates trait covariance and uncertainty, and we map its spatial structure across North American forests. This approach provides a scalable functional complement to species-centred projections by highlighting where communities are most likely to experience novel or suboptimal climate conditions in the coming decades. Following recent work, we selected 18 representative physiological traits spanning leaf economics, wood structure, moisture regulation, belowground allocation, and tree size (Maynard *et al*., 2022; Paz, Crowther and Maynard, 2024; Jimenez *et al*., 2025), in addition to five trait syndromes reflecting abiotic tolerances (Rueda, Godoy and Hawkins, 2018) and information on mycorrhizal symbiosis type (Averill *et al*., 2022). Using forest-inventory data from over 40,000 mature plots (excluding early- and mid-successional stands), we model functional profiles for these 24 traits as a function of climate, soils, and topography. By comparing present-day functional composition with the trait profiles most compatible with projected future climates, we quantify TEM risk for each plot and identify which traits, forest groups, and regions exhibit the lowest and highest risk. We then analyse the abiotic, ecological, and management covariates associated with these patterns. We hypothesise that (1) communities experiencing the most severe drought conditions will show the highest TEM risk and (2) communities with greater species richness will exhibit lower TEM risk due to enhanced functional buffering. Together, this provides the first continental-scale assessment of TEM risk, identifying its drivers and spatial hotspots and highlighting priority locations for monitoring, experimentation, and climate-adapted management.

## Materials and Methods

### Overview

We use a two-step framework to quantify and map trait–environment misalignment risk in North American forests. First, using mature inventory plots from the United States and Canada, we assemble species composition and derive basal-area–weighted community-weighted means (CWMs) for 24 traits (1. Data). We then train a multi-output random forest regressor to predict each plot’s CWM trait vector from its environment, and we use the resulting models to project CWMs under future climate (2. Trait–environment model). By comparing current trait CWMs to future projected CWMs, we identify regions with high vs. low risk of trait-environment misalignment (TEM) (3. Trait–environment misalignment risk calculation). We then map per-plot TEM risk across North America, and we quantify how TEM risk varies across abiotic variables, ecological indicators, and ownership categories (4. Spatial patterns and drivers of TEM risk).

#### 1. Data

##### Forest inventory data

Forest inventory data were sourced from the United States Department of Agriculture (USDA) Forest Inventory and Analysis Program (FIA) and contained plot-level information on stand age, forest group classification, and species composition, as well as tree-level information on stem diameter at breast height (DBH) and trees per acre for all trees > 5 inches DBH (Gray *et al*., 2012). To focus on mature and intact forests, we retained only plots that were fully forested, had living and undamaged trees, and had a stand age in the 50th percentile for their forest group. For consistency, we further limited the analysis to plots surveyed using the FIA’s national plot design or local extensions to this methodology (FIA design codes 1, 111, 112, 113, 116, 117, 311, 312, 501, 502, 503, 504, 505, 506). This resulted in a total dataset of 41,421 US plots. Equivalent information regarding age, size, species, and status for Canadian forest plots was extracted from the Canadian National Forest Inventory (NFI) for the provinces of New Brunswick, Nova Scotia, the Northwest Territories, Ontario, Quebec, and public land (Gillis, Omule and Brierley, 2011). This approach resulted in a final Canadian dataset comprising 886 plots and a total North American dataset of 42,307 plots.

##### Environmental data

We selected two broad types of environmental data: climatic variables shown to play an important role in determining habitat range and trait expression in trees, namely temperature and precipitation means and extremes (Moles *et al*., 2014); and physical properties relating to soil characteristics or topology, such as clay content and elevation, which influence growth, survival, and trait properties (Paulsen, Weber and Körner, 2000; Joswig *et al*., 2022),. This resulted in eight climatic variables: mean annual air temperature, annual air temperature range, mean daily air temperature of the driest quarter, mean daily air temperature of the coldest quarter, annual precipitation, precipitation seasonality, mean monthly precipitation of the driest quarter, and mean monthly precipitation of the coldest quarter. These data were sourced from the CHELSA Bioclim database v2.1 (Karger *et al*., 2017), which provides downscaled model outputs for temperature and precipitation estimates from the ERA-Interim climatic reanalysis at a resolution of 30 arc seconds. Raster layers for each geographic pixel were extracted from GeoTIFF files using the rasterio package in Python (version 1.3) for the period 1981–2010, as well as for 2080–2100 under three Shared Socioeconomic Pathway (SSP) scenarios (SSP126, SSP370, and SSP585), based on projections from the GFDL-ESM4.1 model. The GFDL-ESM4.1 model was chosen due to its high-priority designation under the ISIMIP3b protocol, and the three SSPs represent low, medium, and high ranges of projected climate change. Additionally, current biome information for each plot was extracted from the Terrestrial Ecoregions of the World (Olson *et al*., 2001) shapefile using the sf package (version 1-0-17) in R.

We included 10 physical properties as predictors: four topographical variables—eastness index, northness index (indicating orientation relative to east-west and north-south, respectively, both on a −1 to +1 scale), elevation (metres), and slope (degrees)—were sourced from EarthEnv (Amatulli *et al*., 2018); and six soil physical properties—bulk density (fine earth) at 0.15m, clay content (0–2 μm) at 0.15m, coarse fragment volumetric content at 0.15m, depth to bedrock (up to 200cm), sand content (50–2000 μm) at 0.15m, and silt content (2–50 μm) at 0.15m—were taken from the SoilGrids250m dataset (Hengl *et al*., 2017).

##### Traits and trait syndromes

Species-level tree trait data were taken from three trait databases. First, 18 physiological and morphological traits were taken from Maynard et al., representing leaf economics, wood structure, moisture regulation, belowground allocation, and tree size (Maynard *et al*., 2022). In addition to these physiological traits, we incorporated five trait syndromes given in Rueda et al., including cold tolerance, shade tolerance, drought tolerance, waterlogging tolerance, and fire tolerance (Rueda, Godoy and Hawkins, 2018). These syndromes influence species interactions and habitat requirements (Delavaux *et al*., 2023) and are particularly relevant for silviculture and forest management (Kenefic *et al*., 2021). Although these syndromes ultimately stem from physiological and morphological traits, their relationships with individual traits are relatively weak (Rueda, Godoy and Hawkins, 2018), suggesting that they should be treated as distinct entities when exploring the effects of biotic and abiotic interactions on community structure. Lastly, we extracted information of mycorrhizal symbiosis type from Averill et al., providing insight into belowground allocation and resource uptake which can drive community assembly in North American forests (Averill *et al*., 2022).

The 18 physiological traits were all continuous measurements with imputed data estimated at the species-level using phylogenetic similarity (see Maynard *et al.,* 2022 for details). For four of the six trait syndromes (shade, drought, waterlogging and fire tolerance), species were rated on an ordinal scale (1-4), with higher values indicating greater tolerance. Cold tolerance was represented by the lowest temperature a species can withstand, with the absolute value used to align directionally with other syndromes (higher absolute values indicating greater cold tolerance). For mycorrhizal symbiosis, Averill et al.’s classification was converted into a categorical scale: 1 for arbuscular mycorrhizal (AM), 0 for ectomycorrhizal (ECM), and 0.5 for mixed associations. This gradient value allowed for differentiation between exclusive and mixed types of mycorrhizal associations. Missing trait syndrome data were imputed using a nearest neighbour phylogenetic look-up process based on the V.Phylomaker (Jin and Qian, 2019) package in R (version 0.1.0). The 18 physiological traits and six trait syndromes are hereafter collectively referred to as traits (see Supplementary Information, Supplementary Table S1 for list of 24 traits).

##### Community-weighted trait mean calculation

To quantify each plot’s functional composition, we calculated its community-weighted trait means (CWMs) by first determining the basal area proportion of each tree species per plot. The basal area (BA) per species per plot was calculated using the formula:

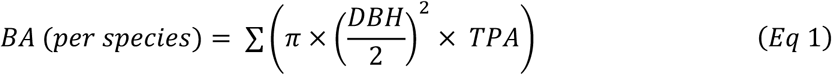

where DBH is the tree diameter at breast height, TPA represents the unadjusted trees per acre (expansion factor), which can vary by plot and condition, and the summation is taken over all individuals of that species within the plot. The relative proportion of species was then taken as the fraction of total basal area within that plot. These proportions were used to derive a community-weighted single mean value for each trait for each forest plot. Traits exhibiting skew (root depth, stem diameter, leaf thickness, crown diameter, tree height, leaf area, bark thickness, seed dry mass and waterlogging tolerance) were log_e_ transformed, and all traits were normalised by removing the mean and scaling to unit variance. Each plot was thus represented by a vector of 24 CWMs.

#### 2. Trait–environment model

We trained a multivariate random forest regressor (RFR) within a multioutput framework (scikit-learn v1.5.2, Python) to simultaneously predict each plot’s vector of CWMs using the environmental variables. Polynomial transformations (squared terms) and pairwise interactions of the 18 present-day environmental variables were generated prior to model fitting and appended as features. The data were split into training (70%) and test (30%) sets, stratified by forest group to ensure balanced representation of community types.

Hyperparameters were selected via an initial cross-validated grid search on a stratified development subset, then fixed for all analyses to avoid leakage in buffered validation. The final RFR was configured with 1,800 trees, a minimum sample split of 2, a minimum of 1 sample per leaf, log2 as the maximum number of features, a maximum depth of 20, and without bootstrapping. These hyperparameters were chosen to balance model complexity and overfitting, with the large number of estimators and controlled depth enabling simultaneous prediction of all 24 traits.

To obtain accurate model validation and to assess the model’s ability to generalise to new conditions, we conducted buffered leave-one-group-out cross-validation (LOGOCV) (Roberts *et al*., 2017) stratified by mean annual surface air temperature and mean annual precipitation (Saxe *et al*., 2001). This approach, implemented using the sklearn model selection package in Python (version 1.5.2), tested the model’s ability to predict CWMs for temperature and precipitation values that had been removed from its training set. For example, if a test group has a mean temperature of 10°C and a 2°C buffer is applied, then all training data with temperatures between 9°C and 11°C are excluded from the model training set. We applied buffered LOGOCV based on the maximum projected increases under the extreme SSP585 scenario. For precipitation, we applied incremental buffers of 40 mm up to 200 mm, and for surface air temperature, we applied buffers of 1°C up to 7°C. R² and normalised RMSE metrics were used to evaluate both the RFR model performance and the buffered LOGOCV.

CWM predictions for each plot were generated for SSP126, SSP370, and SSP585. We primarily report on SSP370 in the main text as it represents a moderate mid-range climate future, while results for SSP126 and SSP585 are provided in the supplementary information. Trait differences were assessed by comparing the normalised predictions for each scenario with present-day CWMs. Paired Wilcoxon Signed-Rank Tests were conducted to analyse the magnitude and significance of trait differences across scenarios using the Scipy stats package in Python (version 1.14.1). A Bonferroni correction was applied to adjust for multiple comparisons, with a significance threshold of p < 0.05.

Sampling uncertainty was assessed using bootstrapped sampling with replacement (n = 500). For each plot and climate scenario, the mean, median, standard deviation (SD), and coefficient of variation (CV) of the predicted CWMs was calculated across the 500 bootstrapped runs.

#### 3. Trait–environment misalignment risk calculation

Trait–environment misalignment (TEM) risk was quantified as the Mahalanobis distance in 24-dimensional CWM space between each plot’s present-day composition and its predicted climate-associated composition under late-century conditions, with larger values indicating greater overall misalignment. Mahalanobis is akin to Euclidean distance but accounts for covariance among traits, minimising the influence of any one trait and reducing inflation of TEM risk due to trait collinearity. Mahalanobis distance is interpretable as a Z-score in units of standard deviation account for correlation among traits.

Each forest plot was assigned to a taxonomic reference group (for example Spruce/fir, Oak/hickory) to provide a common baseline for normalising predicted differences. U.S. plots retained their FIA group classification, while Canadian plots, lacking an equivalent classification, were assigned group labels via a random forest classifier (randomForest library in R, version 4.7-1.1) trained on U.S. data using normalised trait means. These assignments were validated by confirming that species composition in Canadian plots corresponded with their predicted forest group. To demonstrate that plots belonging to the same forest group share similar CWMs, the 24 CWMs per plot were projected into a two-dimensional trait space using the Uniform Manifold Approximation and Projection (UMAP) (McInnes and Healy, 2018) algorithm implemented by the uwot package (Melville, 2019) in R (version 0.2.3; cosine distance; n_neighbors = 5; min_dist = 0.2). This allowed visual verification that plots within the same forest group clustered closely together in two-dimensional trait space.

TEM risk was calculated by first determining the difference between the current and predicted CWMs for each trait in each plot. These differences were then normalised using the forest-group standard deviation (SD) of each trait to account for the natural variability of trait expression within forest groups. Mahalanobis distance was computed in this normalised trait space using a covariance matrix estimated across present-day plots, with numerical regularisation applied to ensure stable inversion under collinearity.

##### Incorporating trait variation

Traits can vary within species due to environmental plasticity and other sources of intraspecific variation, and the species trait values used here were phylogenetically imputed and therefore contain prediction error. To account for both sources of uncertainty, we generated a Monte Carlo ensemble of perturbed present-day CWM vectors for each plot and recalculated TEM risk for each draw. For each physiological trait, perturbations were sampled from empirical relative prediction error distributions for that trait, as given by Maynard et al. (2022). These error distributions were stratified by taxonomic context, using order-level error distributions when sufficiently supported (>30 species within that order) otherwise sampling from broader Angiosperm or Gymnosperm pools. Specifically, for each species-by-trait combination we generated 1000 perturbed present-day trait values, reflecting both trait uncertainty and species-level trait variation. From these, we calculated 1000 perturbed CWM vectors for each plot (Fig. 1, light blue circles) and computed Mahalanobis distance to the same predicted future CWM vector for each draw. The primary TEM risk metric reported was the minimum Mahalanobis distance across the 1000 draws for each plot (Fig. 1, black arrow). This represents the best-case alignment attainable under modelled intraspecific variation and trait-estimation uncertainty, and therefore provides a conservative estimate that deliberately avoids overestimating risk due to measurement and imputation error, while also incorporating the potential for trait acclimation, phenotypic plasticity, or intraspecific variation to offset risk.

#### 4. Spatial patterns and drivers of TEM risk

We summarised spatial patterns of TEM risk by averaging per-plot minimum Mahalanobis distance values at community and regional scales to evaluate trait–environment misalignment across the landscape. Spatial patterns were aggregated within hexagonal pixels to a standard area of 23,322 km² via the dggridR package in R (version 2.0.4).

Additionally, for each trait within each forest group, we quantified trait misalignment as the absolute difference between projected future and current trait values after standardising by the forest group’s current standard deviation for that trait. For each plot, we selected the Monte Carlo replicate that produced the smallest multivariate Mahalanobis distance to the plot’s future trait vector and used the corresponding per-trait misalignment values from that replicate. We then summarised misalignment by averaging these per-trait values within each forest group and visualised the resulting group-by-trait matrix as a heatmap.

##### Trait-environment misalignment risk covariates

To identify covariates associated with future trait–environment misalignment, we fitted a random forest regressor (default parameters) to predict plot-level overall TEM risk from a hypothesis-driven set of environmental, ecological, and ownership-related covariates.

Predictors comprised climate (mean annual air temperature, annual temperature range, mean daily mean temperature of the coldest quarter, annual precipitation, precipitation seasonality, mean monthly precipitation of the driest quarter), climate change (change in mean annual temperature and annual precipitation), topography (elevation, slope), soils (clay content, depth to bedrock), ecology (species richness, stand age), and ownership factors (human activity, ownership group). Owing to gaps in Canadian covariate coverage, analyses were restricted to U.S. plots. We derived SHapley Additive exPlanations (Shapley, 1953; Lundberg and Lee, 2017) to assess feature importance and summarised results by biome to identify broader ecological patterns. SHAP was used because it quantifies, for each plot, how much each predictor increases or decreases modelled TEM risk, providing a clearer and more comparable measure across biomes than native feature-importance scores. To show how each covariate relates to its SHAP contribution, we plotted SHAP dependence scatterplots for a subset of predictors (number of species, stand age, precipitation seasonality, precipitation, human activity) and overlaid a Generalised Additive Model (GAM) line of best fit with 95% confidence shading to summarise any non-linear patterns.

## Results

### Differences in functional trait envelopes under climate change

The random forest models had an average cross-validated R² value of 0.61 across all traits, and a normalised root mean square error (NRMSE) of 0.04 (see Supplementary Information, Supplementary Fig. S2). The buffered Leave-One-Group-Out Cross-Validation (LOGOCV) across the ranges of climatic values likely to be realised under the SSP scenarios demonstrated good retention of predictive accuracy, with the R² remaining above 0.50 and NRMSE at 0.08 for mean annual precipitation, even when predicting SSP585 levels of precipitation change (Supplementary Fig. S3a). For surface air temperature, R² decreased to 0.37 for SSP370 changes and to 0.29 for SSP585, while NRMSE remained consistent at 0.07 and 0.08, respectively (Supplementary Fig. S3b).

For SSP370, the largest differences in functional envelopes were predicted for the tolerances rather than physiological traits. Cold tolerance values showed the greatest projected decline (mean scaled difference: −0.218, p < 0.0001), followed by a significant decrease in shade tolerance (−0.151, p < 0.0001). In contrast, drought tolerance values were forecast to increase substantially (+0.116, p < 0.0001). The single largest increase in trait values was predicted for the reproductive trait, seed dry mass (+0.186, p < 0.0001). Among structural traits, tree height was projected to decrease markedly (−0.142, p < 0.0001), alongside reductions in crown height (−0.038, p < 0.0001) and stem diameter (−0.036, p < 0.0001) (Fig. 3a).

**Figure 3:**
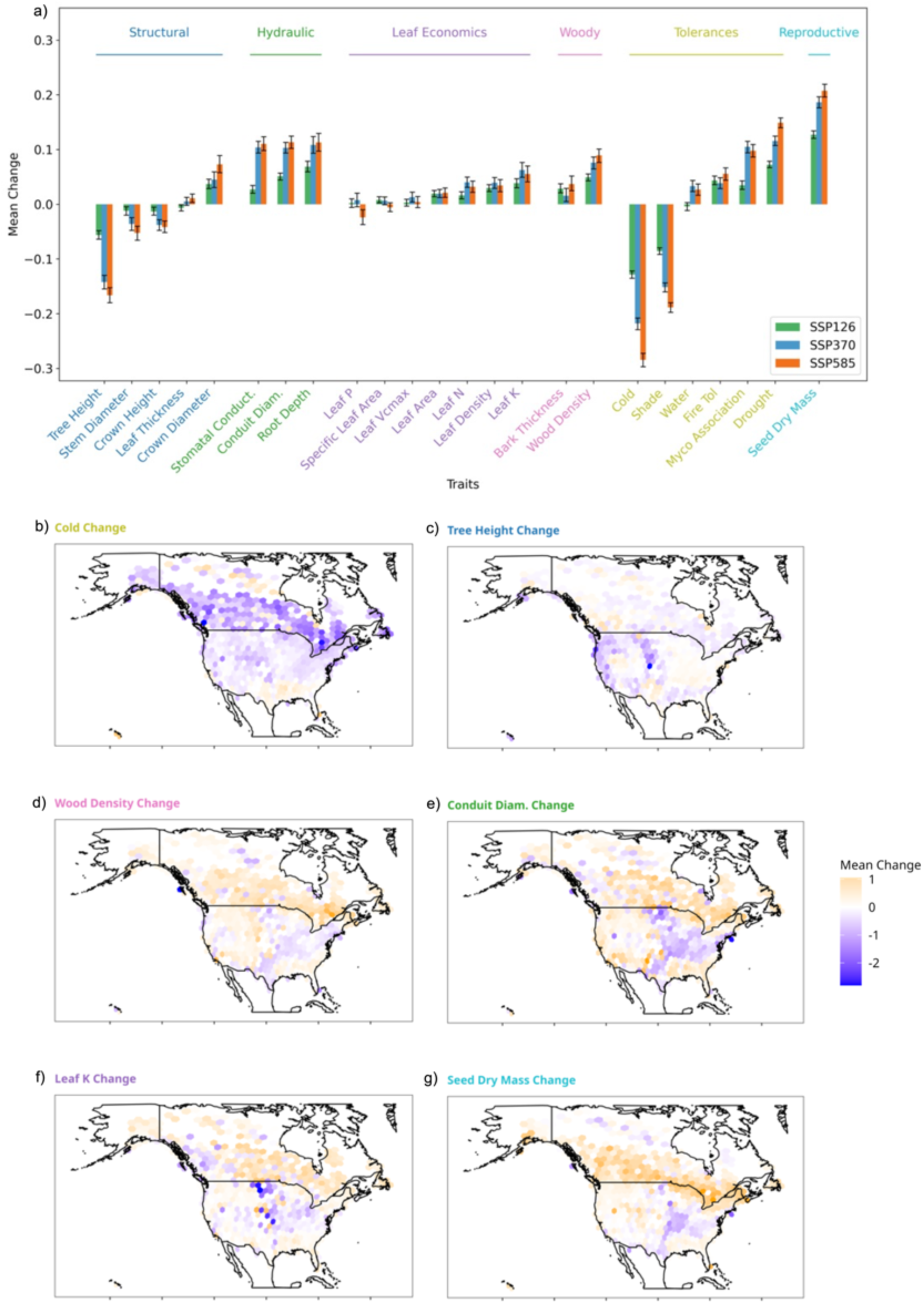
Mean differences in compatible community trait means and spatial distribution of differences. a) The x-axis represents traits, and the y-axis shows the mean predicted normalised difference in scaled community-weighted trait means across all plots relative to current values after 500 bootstrapped model runs (sampling with replacement). Traits grouped into categories – structural, hydraulic, leaf economics, woody, tolerances and reproductive - and coloured. Whiskers indicate the 5% and 95% empirical bootstrap confidence intervals, while columns represent different SSP scenarios. b-g) Spatial distribution of largest scaled trait differences in each category.

Across North America, larger hydraulic trait values were predicted to be more compatible with future climate, on average, including higher stomatal conductance (mean scaled difference: +0.104, p < 0.0001), root depth (+0.108, p < 0.0001), and conduit diameter (+0.103, p < 0.0001). Woody structural traits also showed significant differences between current composition and future compatible functional envelopes, with higher wood density being moderately more favourable (+0.076, p < 0.0001), while bark thickness had a projected smaller but significant increase (+0.016, p = 0.00039). Leaf economics traits showed mixed predicted responses. Leaf nitrogen (+0.04, p < 0.0001) and leaf potassium (+0.063, p < 0.0001) concentrations were forecast to increase, while leaf phosphorus (+0.008, p < 0.0001) had minimal differences. Leaf density was predicted to increase (+0.039, p < 0.0001), while specific leaf area (SLA) had a projected slight but significant increase (+0.006, p < 0.0001) (Figure 3). See Supplementary Table S1 for trait differences and Table S2 for Wilcoxon Signed-Rank trait change test results.

Projected trait differences exhibited clear spatial patterns. The most pronounced declines in cold (Fig. 3b) and shade tolerance appeared at higher latitudes (∼60° N), alongside reductions in tree height (Fig. 3c), leaf thickness and crown height, and increases in wood density (Fig. 3d). In these regions, hydraulic traits—including conduit diameter (Fig. 3e) and stomatal conductance—and leaf economic traits (leaf potassium (Fig. 3f), nitrogen and phosphorus concentrations, SLA and seed dry mass (Fig. 3g)) showed notable increases. By contrast, much of the mid-eastern and north-eastern United States displayed the opposite trend: projected rises in drought tolerance and structural traits (stem diameter), and woody traits (wood density (Fig. 3d), bark thickness), accompanied by declines in hydraulic traits (conduit diameter (Fig. 3e)) and leaf economic traits—leaf potassium (Fig. 3f), nitrogen and phosphorus concentrations, SLA—as well as seed dry mass (Fig. 3g). See Supplementary Figure S4 for spatial projections of all trait differences.

### Spatial patterns in trait-environment misalignment risk

Elevated trait–environment misalignment risk was predicted for western and central North America, concentrated in mountain conifer forests and in prairie regions extending from the central United States into Canada (Figure 4a). The northern Rocky Mountains—across British Columbia, Alberta, Montana, Idaho, and Wyoming—contained extensive areas of higher risk (TEM risk > 5 standard deviations (SDs)), with some locations around 7–8 SDs. In these montane forests, the highest-risk areas were linked mainly to misalignments in water-use and leaf-function traits and were most common in forest groups such as Lodgepole pine and Spruce / fir (Supplementary Information, Supplementary Figure S5).

**Figure 4:**
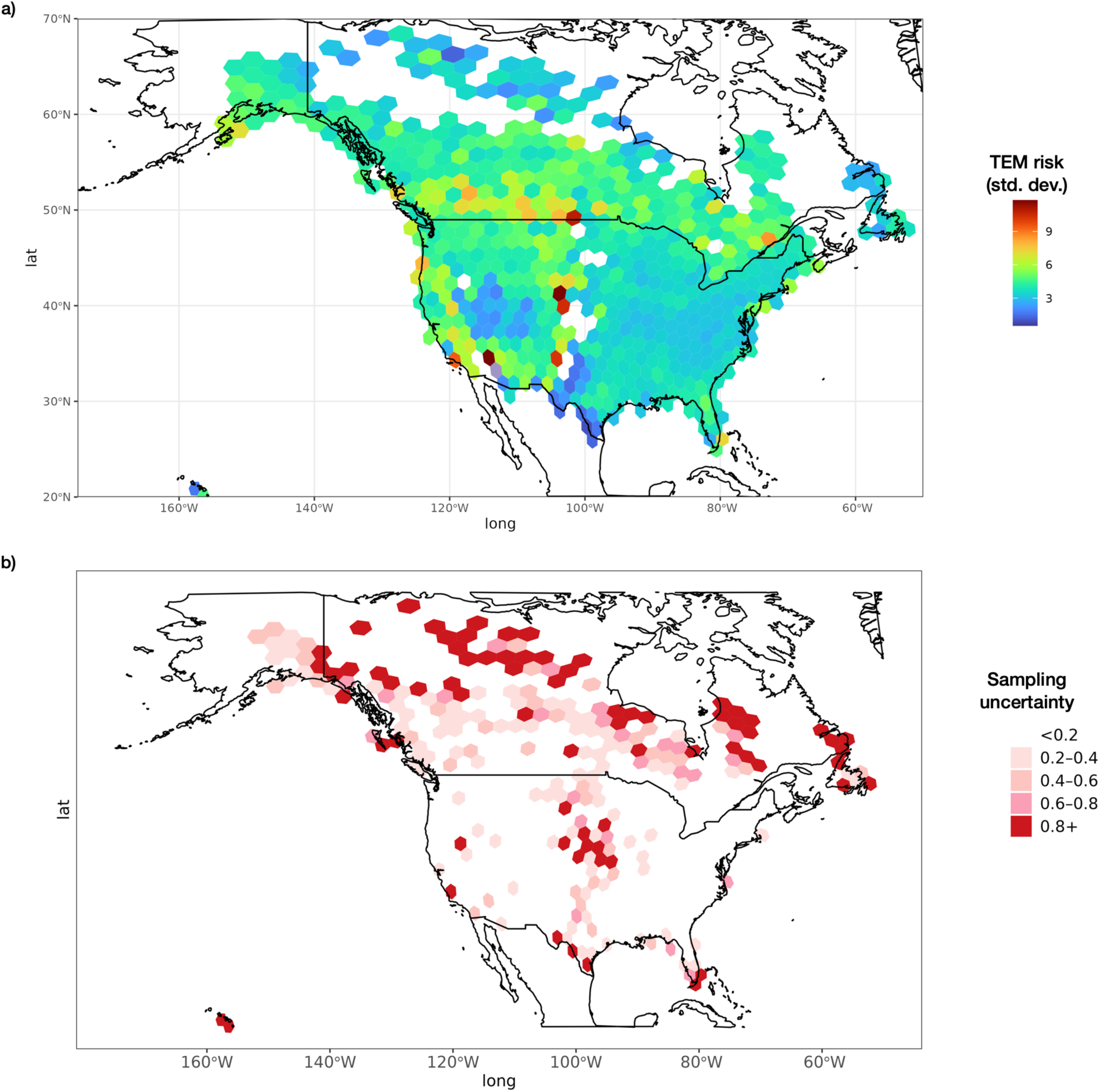
Spatial projections of trait–environment misalignment risk. (a) Risk of trait-environment mismatch (minimum Mahalanobis distance) in 24-trait CWM space between present-day and 2100 climate-associated composition (higher values indicate greater overall misalignment) taken from 1000 Monte Carlo draws. Units are standard deviations of Mahalanobis distance. (b) Sampling uncertainty, shown as the coefficient of variation in risk across 500 bootstrap replicates.

High TEM risk was also projected for semi-arid landscapes, including the western Great Plains and intermontane basins. High-risk areas were concentrated in eastern Montana and Wyoming, the Dakotas, and north into southern Alberta and Saskatchewan (Figure 4a). Within the sparser prairie–forest transition, hotspots reached about 10–11 SDs, indicating very strong predicted misalignment in parts of the Canadian Prairies and the adjacent northern Great Plains. In these dry-edge regions, misalignment was strongest for traits linked to woody growth and to responses to fire and drought, with high-risk groups including Ponderosa pine and White / red / jack pine (Supplementary Information, Supplementary Figure S5).

Coastal and inland regions of California also had elevated TEM risk and moderate-to-high risk extended across both coastal and inland forests of the Pacific Northwest (Figure 4a). Along the Pacific coast, the highest-risk forests included Redwood, where misalignment was concentrated in leaf-function and water-use traits. Farther inland in the Pacific Northwest, misalignment risk reflected a broader mix of water-use and structural traits, with affected groups including Douglas-fir and Hemlock / Sitka spruce (Supplementary Information, Supplementary Figure S5).

In contrast, the northeastern United States had lower predicted TEM risk overall than the western and prairie regions, with values most often around 2–4 SDs across the Appalachian region and Atlantic coastal lowlands (Figure 4a). These lower-risk regions were associated with FIA groups that showed smaller overall misalignments, including broadleaf-dominated assemblages such as Oak / hickory, Oak / pine, and Western oak. Xeric and desert regions containing Woodland hardwoods in the southern United States (Texas) had the lowest predicted risk in the study area, generally below 3 SDs and in some locations around 1.5 SDs (Figure 4a).

Areas with low forest cover at ecological margins—such as high-latitude tundra regions and grassland–savanna ecoregions in the Great Plains—combined high TEM risk (TEM risk > 5 SDs) with higher uncertainty (high CV) (Figure 4b). Uncertainty was low for most predictions (CV < 0.2 in 90% of cases), and higher uncertainty was concentrated near ecological margins and in regions with sparse forest cover (Figure 4b). Spatial patterns of projected trait differences and risk of trait-environment misalignment were broadly consistent across all sensitivity analyses, irrespective of whether calculations used mean trait difference (Supplementary Figure S6), mean or median TEM risk (Supplementary Figures S7, S8). TEM was shown to have a broadly monotonic relationship with severity of SSP, with the highest levels of TEM observed under SSP585 (Supplementary Figure S9).

### Drivers of trait-environment misalignment risk

Trait-environment misalignment risk values were predominantly influenced by species richness (SHAP value: 0.409), precipitation seasonality (0.268) and stand age (0.111) (Fig. 5a). A strong negative relationship was observed between species richness and TEM risk, with a sharp decline in risk as richness increased from monocultures to seven-species systems, followed by a more gradual decrease up to ten species and minimal additional effect beyond (Fig. 5b). Forests experiencing higher precipitation seasonality were associated with higher TEM risk, especially in temperate conifer biomes (Fig. 5c). Stand age exhibited mixed trends, with a general pattern of greater stand age corresponding to higher TEM risk (Fig. 5d). Changes in mean annual temperature displayed no clear overall trend (Fig. 5e). Human activity had a low overall SHAP value (0.009) but exhibited a clear pattern, with forests that had experienced human activity showing higher TEM risk than undisturbed forests (Fig. 5f).

**Figure 5:**
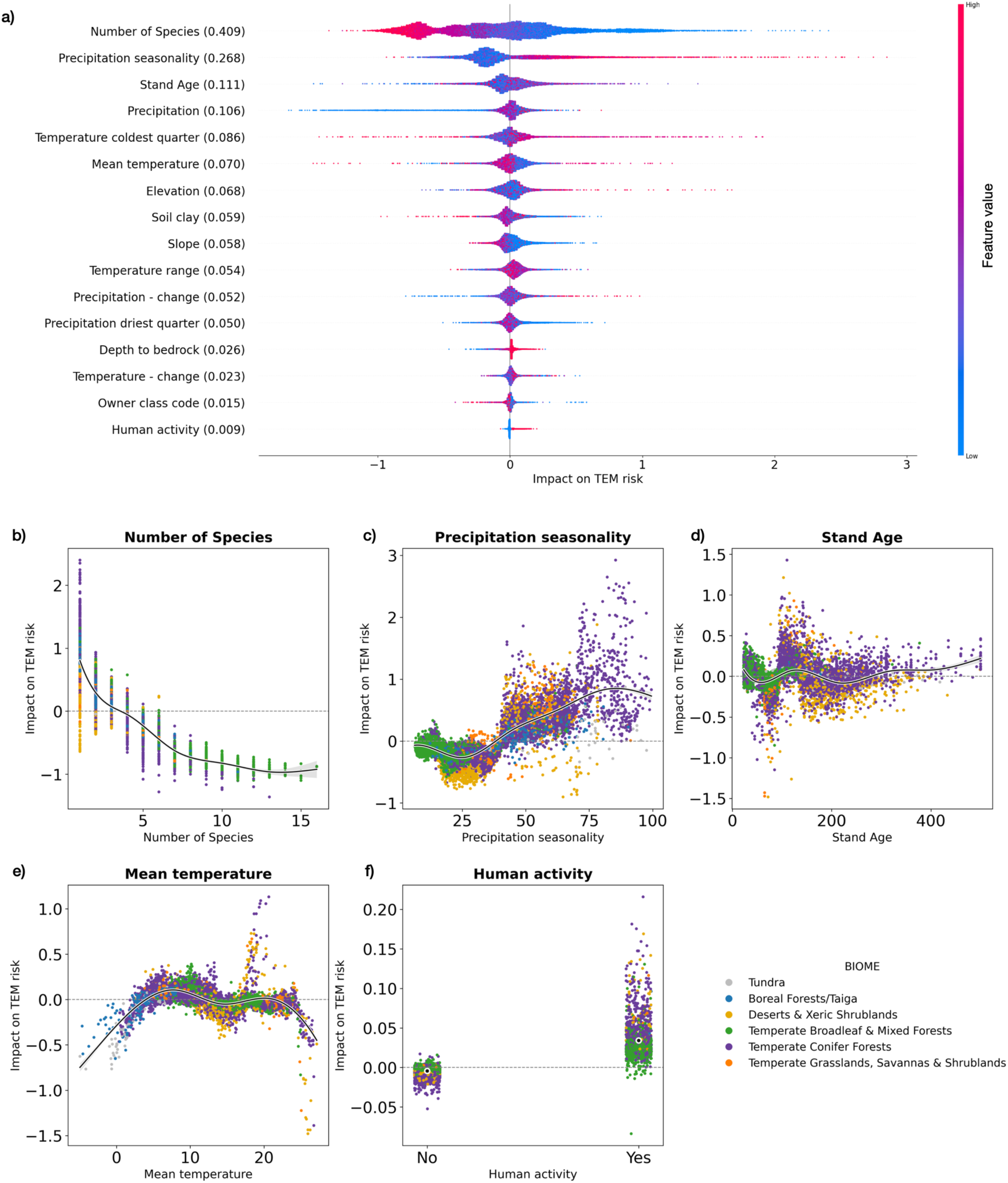
Covariates of trait-environment misalignment risk. (a) Each point represents a forest plot, with positive x-axis values indicating an increase in TEM risk and negative values indicating a decrease. Higher covariate values are shown in red, lower in blue. The overall absolute risk impact value for each covariate is displayed in brackets. For example, plots with higher numbers of species (points clustered in red) are associated with lower predicted risk (positioned further to the left on the x-axis). (b–f) risk value distributions for individual covariates. The x-axis represents covariate values, the y-axis shows impact on TEM risk values, and each point corresponds to a forest plot. Colour represents biome type.

## Discussion

Our results indicate that projected climate change—including reduced cold limitation at higher latitudes and increased drought stress in the eastern United States (Supplementary Figure S1)—is likely to push many forests into climate conditions beyond those associated with their current trait profiles. However, despite widespread projected differences in trait values, predicted trait–environment misalignment risk is strongly spatially structured, with the highest values concentrated at high latitudes and in montane conifer forests. In contrast, forests with greater species richness show lower risk, suggesting that local diversity can buffer functional misalignment under future climate change.

The goal of our analysis is to identify which forests are most at risk of being incompatible with projected climates, not to make temporal projections of future functional composition nor forecast “optimal” assemblages. We therefore stress that higher TEM risk is an indicator that communities are more *likely* to experience functional misalignment under projected climate conditions, but that there are numerous ecological and evolutionary reasons why this risk may not actually be realised. For example, local adaptation and acclimation, together with phenotypic and ontogenetic plasticity, may allow existing trees and forest communities to shift their trait profiles in line with a changing climate (Leites and Benito Garzón, 2023). Indeed, here we explicitly address this by incorporating intraspecific trait variation into our modelling framework to provide conservative estimates that account for the range of plausible traits. As expected, if we did not account for trait variation, then TEM risk values would be slightly higher (Supplementary Fig S10), yet the spatial patterns of misalignment remain consistent, highlighting that trait variation can reduce risk, but is unlikely to offset these risks altogether. Alternatively, forest biogeographic patterns may not be in climatic equilibrium, such that some forest types (and functional profiles) may have substantially wider climate envelopes than observed in the data (Laughlin and McGill, 2024). For such forest types, our results can therefore be interpreted as an upper bound on risk. Lastly, mature adult trees are often able to withstand abiotic conditions that are unfavourable to developing trees, facilitating the persistence of forest types that would otherwise be unable to regenerate under future climate (Svenning and Sandel, 2013). Given that our projections extend approximately 75 years into the future (2100)—far less than the life expectancy of many trees—this risk is therefore more likely to manifest through its effects on the functional capacity and health of existing trees, rather than through community turnover per se.

Should climate-driven functional misalignment risk be realised in specific forests, the consequences are often expected to be detrimental. Reported consequences include increased physiological stress (O’Brien *et al*., 2017; Quetin *et al*., 2023), growth declines and productivity losses (Morin *et al*., 2018; Ammer, 2019), higher mortality (Allen *et al*., 2010; Anderegg *et al*., 2016), stronger impacts of fire, insects and pathogens (Seidl *et al*., 2017; Oliva, Redondo and Stenlid, 2020), and greater likelihood of community transition after extremes (Johnstone *et al*., 2016; Batllori *et al*., 2020). Yet positive consequences may also occur, particularly in cold-limited systems where warming lengthens growing seasons and can raise productivity over limited ranges and time windows (Way and Oren, 2010). By identifying regions likely to face the largest trait–environment misalignment, this work can help prioritise research into when and how adjustment occurs and what consequences follow. Regions with high misalignment risk, for example, could be prioritised for field-based survey work and plot remeasurement to track trajectories (Michalak *et al*., 2022), evaluate the role of local adaptation and phenotypic plasticity, inform manipulations of composition to test underlying mechanisms (Nagel *et al*., 2017), and guide climate-smart trials to evaluate management options (Bower *et al*., 2024).

Contrary to our hypothesis that drought-affected communities would show the highest misalignment risk, many of these communities actually exhibited relatively low risk of misalignment. In mid-latitude and north-eastern regions, our model shows rising drought stress would favour traits linked to water conservation, with narrower conduits, lower SLA, and reduced foliar nutrients indicating a shift towards survival and slower growth with increased drought tolerance (de L. Dantas, Batalha and Pausas, 2013; Maracahipes *et al*., 2018). In many cases, the predicted trait differences largely extend existing functional strategies, suggesting that less functional adjustment is required than we hypothesised. Drought-adapted, temperate broadleaf and mixed groups such as Oak/pine forests (Cavender-Bares, 2019) in the eastern and southeastern U.S. exhibit lower trait–environment misalignment risk, as their existing trait profiles are closer to the climate-associated trait profiles expected under projected conditions.

In contrast, our results project that milder conditions in high latitude regions at the transition between temperate and boreal systems may reduce the advantage of cold and shade tolerance, favouring increased hydraulic efficiency and carbon assimilation through wider conduits (McCulloh *et al*., 2016), deeper roots (Pregitzer *et al*., 2000), and higher stomatal conductance (Urban *et al*., 2017). These shifts would favour more acquisitive growth strategies, as indicated by higher foliar nitrogen, potassium, and phosphorus (Maracahipes *et al*., 2018). However, this would come at the cost of reductions in leaf thickness, crown height, and stem diameter, suggesting a move away from cold-adapted stress tolerance towards shorter-statured, faster-growing communities (Maracahipes *et al*., 2018). For cold-adapted gymnosperm forests such as Lodgepole pine, Spruce/fir, and White/red/jack pine these shifts correspond to higher trait–environment misalignment risk and therefore imply a larger functional adjustment relative to the trait profiles typically associated with projected conditions. In the boreal transition zone (British Columbia, Yukon, and Alaska) and the northern Rocky Mountains, where these groups are dominant, their reliance on conservative hydraulic strategies may constrain adjustment under increasing warmth, with potential consequences for competitive dynamics, composition, and carbon storage (Carnicer *et al*., 2013; Dad, Rashid and Chen, 2023).

Consistent with our hypothesis, higher species richness was associated with lower trait–environment misalignment risk across all groups, supporting a positive role for diversity in buffering community-level functional change. Communities with greater richness showed lower TEM risk, plausibly because a larger species pool expands the range of functional traits present, reducing the chance that late-century climatic shifts push communities beyond their existing functional envelope (Dolezal *et al*., 2024). This pattern aligns with the diversity–stability framework, in which richer communities are stabilised by redundancy and compensatory dynamics that help maintain functioning under environmental variability (McCann, 2000). This buffering does not require wholesale turnover, since even modest increases in richness can introduce functionally distinct taxa that widen realised trait space and reduce the probability of strong misalignment under projected conditions (Hooper *et al*., 2005). Richness can also promote stability through partial asynchrony in species’ responses to climate variability, damping temporal swings in community-level properties as different taxa respond differently to anomalies (Ives and Carpenter, 2007). Our results provide additional support for these reported mechanisms and suggest that their stabilising effects extend to maintaining trait–environment alignment under projected late-century climates.

This work provide the first continental-scale, functional assessment of forests’ risk of misalignment with future environmental conditions, but as with any large-scale trait analyses, these results come with important considerations (Wilkes *et al*., 2020). Spatial biases in global datasets can complicate efforts to interpolate patterns beyond the observed data. While the FIA sampling design provides systematic coverage of U.S. forests, differences in data density across the US and Canada could still contribute to regional variation in prediction confidence (i.e., higher uncertainty in high latitude regions). In addition, although our focus is on mature forests, stand age was a significant driver of TEM risk (Fig. 5a), and the incorporation of successional dynamics and disturbance regimes could help distinguish transient functional changes from permanent shifts in composition, and thereby refine long-term projections (Meiners *et al*., 2015; Wang *et al*., 2015; Larocque *et al*., 2024).

Ultimately, the extent to which communities experience disruption will depend on the interaction between the magnitude and direction of trait shifts and the community’s ability to buffer change. An important outstanding question is the extent to which TEM risk will align with compositional shifts, which is critical for identifying the disappearance of rare and unique forest types, or the emergence of functionally and compositionally novel communities (Gougherty *et al*., 2024). Linking trait–environment misalignment risk to downstream ecosystem processes—such as carbon sequestration, biodiversity maintenance, and timber productivity—represents an important next step for translating these projections into ecological and socio-economic priorities, and for guiding conservation and climate-adapted forest management (Magalhães Filho *et al*., 2021).

## Conclusion

Our findings indicate widespread shifts in the functional envelopes of North American forest communities, with temperate and montane coniferous ecosystems showing high risk of trait–environment misalignment under projected climate change. However, risk of trait-environment misalignment is not uniform and depends on the interaction between the magnitude and direction of trait differences and a community’s capacity to accommodate those shifts. Where projected differences reinforce existing strategies, misalignment risk is low, but where they oppose incumbent strategies and exceed the range represented within current trait envelopes, risk of misalignment increases sharply. These findings emphasise the need to move beyond species-based assessments of risk and consider the broader functional landscape, with implications for conservation and management strategies aimed at maintaining ecosystem stability under future climates.

## References

Allen, C.D. et al. (2010) ‘A global overview of drought and heat-induced tree mortality reveals emerging climate change risks for forests’, Forest Ecology and Management, 259(4), pp. 660–684. Available at: 10.1016/j.foreco.2009.09.001.

Amatulli, G. et al. (2018) ‘A suite of global, cross-scale topographic variables for environmental and biodiversity modeling’, Scientific Data, 5(1), p. 180040. Available at: 10.1038/sdata.2018.40.

Ammer, C. (2019) ‘Diversity and forest productivity in a changing climate’, New Phytologist, 221(1), pp. 50–66. Available at: 10.1111/nph.15263.

Anderegg, W.R.L. et al. (2016) ‘Meta-analysis reveals that hydraulic traits explain cross-species patterns of drought-induced tree mortality across the globe’, Proceedings of the National Academy of Sciences, 113(18), pp. 5024–5029. Available at: 10.1073/pnas.1525678113.

Andrew, S.C. et al. (2022) ‘Assessing the vulnerability of plant functional trait strategies to climate change’, Global Ecology and Biogeography, 31(6), pp. 1194–1206. Available at: 10.1111/geb.13501.

Averill, C. et al. (2022) ‘Alternative stable states of the forest mycobiome are maintained through positive feedbacks’, Nature Ecology & Evolution, 6(4), pp. 375–382. Available at: 10.1038/s41559-022-01663-9.

Batllori, E. et al. (2020) ‘Forest and woodland replacement patterns following drought-related mortality’, Proceedings of the National Academy of Sciences, 117(47), pp. 29720–29729. Available at: 10.1073/pnas.2002314117.

Bower, A.D. et al. (2024) ‘A practical framework for applied forestry assisted migration’, Frontiers in Forests and Global Change, 7. Available at: 10.3389/ffgc.2024.1454329.

Cadotte, M.W., Carscadden, K. and Mirotchnick, N. (2011) ‘Beyond species: functional diversity and the maintenance of ecological processes and services’, Journal of Applied Ecology, 48(5), pp. 1079–1087. Available at: 10.1111/j.1365-2664.2011.02048.x.

Campbell, E.T. and Brown, M.T. (2012) ‘Environmental accounting of natural capital and ecosystem services for the US National Forest System’, *Environment*, Development and Sustainability, 14(5), pp. 691–724. Available at: 10.1007/s10668-012-9348-6.

Carnicer, J. et al. (2013) ‘Contrasting trait syndromes in angiosperms and conifers are associated with different responses of tree growth to temperature on a large scale’, Frontiers in Plant Science, 4. Available at: 10.3389/fpls.2013.00409.

Cavender-Bares, J. (2019) ‘Diversification, adaptation, and community assembly of the American oaks (Quercus), a model clade for integrating ecology and evolution’, New Phytologist, 221(2), pp. 669–692. Available at: 10.1111/nph.15450.

Dad, J.M., Rashid, I. and Chen, A. (2023) ‘Is climate change pushing gymnosperms against the wall in the northwestern Himalayas?’, Regional Environmental Change, 23(2), p. 51. Available at: 10.1007/s10113-023-02050-1.

Delavaux, C.S. et al. (2023) ‘Mycorrhizal feedbacks influence global forest structure and diversity’, Communications Biology, 6(1), pp. 1–11. Available at: 10.1038/s42003-023-05410-z.

Dolezal, J. et al. (2024) ‘Diversity effects and compensatory dynamics drive productivity and stability in temperate old-growth forests’, Journal of Ecology, 112(10), pp. 2249–2263. Available at: 10.1111/1365-2745.14391.

Enquist, B.J., et al. (2015) ‘Chapter Nine - Scaling from Traits to Ecosystems: Developing a General Trait Driver Theory via Integrating Trait-Based and Metabolic Scaling Theories’, in S. Pawar, G. Woodward, and A.I. Dell (eds) Advances in Ecological Research. Academic Press (Trait-Based Ecology - From Structure to Function), pp. 249–318. Available at: 10.1016/bs.aecr.2015.02.001.

FAO (2022) *The state of the World’s forests* 2022. Available at: 10.4060/cb9360en.

Gillis, M.D., Omule, A.Y. and Brierley, T. (2011) ‘Monitoring Canada’s forests: The National Forest Inventory’, The Forestry Chronicle [Preprint]. Available at: 10.5558/tfc81214-2.

Gougherty, A.V. et al. (2024) ‘Climate Change and the Emergence of No-Analog Forest Assemblages in North America’, Global Change Biology, 30(12), p. e17605. Available at: 10.1111/gcb.17605.

Gray, A. et al. (2012) ‘Forest Inventory and Analysis Database of the United States of America (FIA)’, Biodiversity & Ecology, 4, pp. 225–231. Available at: 10.7809/b-e.00079.

Green, S.J. et al. (2022) ‘Trait-based approaches to global change ecology: moving from description to prediction’, Proceedings of the Royal Society B: Biological Sciences, 289(1971), p. 20220071. Available at: 10.1098/rspb.2022.0071.

Hengl, T. et al. (2017) ‘SoilGrids250m: Global gridded soil information based on machine learning’, PLOS ONE, 12(2), p. e0169748. Available at: 10.1371/journal.pone.0169748.

Herweijer, C. et al. (2020) ‘Nature risk rising: Why the crisis engulfing nature matters for business and the economy’, World Economic Forum and PwC. http://www3.weforum.org/docs/WEF_New_Nature_Economy_Report_2020.pdf.

Hevia, V. et al. (2017) ‘Trait-based approaches to analyze links between the drivers of change and ecosystem services: Synthesizing existing evidence and future challenges’, Ecology and Evolution, 7(3), pp. 831–844. Available at: 10.1002/ece3.2692.

Hooper, D.U. et al. (2005) ‘Effects of Biodiversity on Ecosystem Functioning: A Consensus of Current Knowledge’, Ecological Monographs, 75(1), pp. 3–35. Available at: 10.1890/04-0922.

Ives, A.R. and Carpenter, S.R. (2007) ‘Stability and Diversity of Ecosystems’, Science, 317(5834), pp. 58–62. Available at: 10.1126/science.1133258.

Jimenez, Y.G. et al. (2025) ‘A Global Regionalisation of Tree Functional Capacity’, Global Ecology and Biogeography, 34(7), p. e70083. Available at: 10.1111/geb.70083.

Jin, Y. and Qian, H. (2019) ‘V.PhyloMaker: an R package that can generate very large phylogenies for vascular plants’, Ecography, 42(8), pp. 1353–1359. Available at: 10.1111/ecog.04434.

Johnstone, J.F. et al. (2016) ‘Changing disturbance regimes, ecological memory, and forest resilience’, Frontiers in Ecology and the Environment, 14(7), pp. 369–378. Available at: 10.1002/fee.1311.

Joswig, J.S. et al. (2022) ‘Climatic and soil factors explain the two-dimensional spectrum of global plant trait variation’, Nature Ecology & Evolution, 6(1), pp. 36–50. Available at: 10.1038/s41559-021-01616-8.

Kandlikar, G.S., Kleinhesselink, A.R. and Kraft, N.J.B. (2022) ‘Functional traits predict species responses to environmental variation in a California grassland annual plant community’, Journal of Ecology, 110(4), pp. 833–844. Available at: 10.1111/1365-2745.13845.

Karger, D.N. et al. (2017) ‘Climatologies at high resolution for the earth’s land surface areas’, Scientific Data, 4(1), p. 170122. Available at: 10.1038/sdata.2017.122.

Keenan, R.J. (2015) ‘Climate change impacts and adaptation in forest management: a review’, Annals of Forest Science, 72(2), pp. 145–167. Available at: 10.1007/s13595-014-0446-5.

Kenefic, L.S. et al. (2021) ‘Mixedwood silviculture in North America: the science and art of managing for complex, multi-species temperate forests’, Canadian Journal of Forest Research, 51(7), pp. 921–934. Available at: 10.1139/cjfr-2020-0410.

de L. Dantas, V., Batalha, M.A. and Pausas, J.G. (2013) ‘Fire drives functional thresholds on the savanna–forest transition’, Ecology, 94(11), pp. 2454–2463. Available at: 10.1890/12-1629.1.

Lajoie, G. and Vellend, M. (2018) ‘Characterizing the contribution of plasticity and genetic differentiation to community-level trait responses to environmental change’, Ecology and Evolution, 8(8), pp. 3895–3907. Available at: 10.1002/ece3.3947.

Lamarque, P. et al. (2014) ‘Plant trait-based models identify direct and indirect effects of climate change on bundles of grassland ecosystem services’, Proceedings of the National Academy of Sciences, 111(38), pp. 13751–13756. Available at: 10.1073/pnas.1216051111.

Larocque, G.R. et al. (2024) ‘Simulating the Long-Term Response of Forest Succession to Climate Change in the Boreal Forest of Northern Ontario, Canada’, Forests, 15(8), p. 1417. Available at: 10.3390/f15081417.

Laughlin, D.C. and McGill, B.J. (2024) ‘Trees have overlapping potential niches that extend beyond their realized niches’, Science, 385(6704), pp. 75–80. Available at: 10.1126/science.adm8671.

Leites, L. and Benito Garzón, M. (2023) ‘Forest tree species adaptation to climate across biomes: Building on the legacy of ecological genetics to anticipate responses to climate change’, Global Change Biology, 29(17), pp. 4711–4730. Available at: 10.1111/gcb.16711.

Lindner, M. et al. (2014) ‘Climate change and European forests: What do we know, what are the uncertainties, and what are the implications for forest management?’, Journal of Environmental Management, 146, pp. 69–83. Available at: 10.1016/j.jenvman.2014.07.030.

Lundberg, S.M. and Lee, S.-I. (2017) ‘A Unified Approach to Interpreting Model Predictions’, Advances in Neural Information Processing Systems. Curran Associates, Inc. Available at: https://proceedings.neurips.cc/paper_files/paper/2017/hash/8a20a8621978632d76c43dfd28b67767-Abstract.html (Accessed: 25 February 2026).

Magalhães Filho, L., et al. (2021) ‘A Global Meta-Analysis for Estimating Local Ecosystem Service Value Functions’, Environments, 8(8), p. 76. Available at: 10.3390/environments8080076.

Maracahipes, L. et al. (2018) ‘How to live in contrasting habitats? Acquisitive and conservative strategies emerge at inter- and intraspecific levels in savanna and forest woody plants’, Perspectives in Plant Ecology, Evolution and Systematics, 34, pp. 17–25. Available at: 10.1016/j.ppees.2018.07.006.

Maynard, D.S. et al. (2022) ‘Global relationships in tree functional traits’, Nature Communications, 13(1), p. 3185. Available at: 10.1038/s41467-022-30888-2.

McCann, K.S. (2000) ‘The diversity–stability debate’, Nature, 405(6783), pp. 228–233. Available at: 10.1038/35012234.

McCulloh, K.A. et al. (2016) ‘Is it getting hot in here? Adjustment of hydraulic parameters in six boreal and temperate tree species after 5 years of warming’, Global Change Biology, 22(12), pp. 4124–4133. Available at: 10.1111/gcb.13323.

McInnes, L. and Healy, J. (2018) ‘UMAP: Uniform Manifold Approximation and Projection for Dimension Reduction’, arXiv: Machine Learning [Preprint].

Meiners, S.J. et al. (2015) ‘Is successional research nearing its climax? New approaches for understanding dynamic communities’, Functional Ecology, 29(2), pp. 154–164. Available at: 10.1111/1365-2435.12391.

Melville, J. (2019) ‘uwot: The Uniform Manifold Approximation and Projection (UMAP) Method for Dimensionality Reduction’. Available at: 10.32614/CRAN.package.uwot.

Michalak, J.L. et al. (2022) ‘Climate-change vulnerability assessments of natural resources in U.S. National Parks’, Conservation Science and Practice, 4(7), p. e12703. Available at: 10.1111/csp2.12703.

Michaletz, S.T. et al. (2014) ‘Convergence of terrestrial plant production across global climate gradients’, Nature, 512(7512), pp. 39–43. Available at: 10.1038/nature13470.

Mildrexler, D. et al. (2016) ‘A forest vulnerability index based on drought and high temperatures’, Remote Sensing of Environment, 173, pp. 314–325. Available at: 10.1016/j.rse.2015.11.024.

Miller, J.E.D., Damschen, E.I. and Ives, A.R. (2019) ‘Functional traits and community composition: A comparison among community-weighted means, weighted correlations, and multilevel models’, Methods in Ecology and Evolution, 10(3), pp. 415–425. Available at: 10.1111/2041-210X.13119.

Moles, A.T. et al. (2014) ‘Which is a better predictor of plant traits: temperature or precipitation?’, Journal of Vegetation Science, 25(5), pp. 1167–1180. Available at: 10.1111/jvs.12190.

Morin, X. et al. (2018) ‘Long-term response of forest productivity to climate change is mostly driven by change in tree species composition’, Scientific Reports, 8(1), p. 5627. Available at: 10.1038/s41598-018-23763-y.

Muscarella, R. and Uriarte, M. (2016) ‘Do community-weighted mean functional traits reflect optimal strategies?’, Proceedings of the Royal Society B: Biological Sciences, 283(1827), p. 20152434. Available at: 10.1098/rspb.2015.2434.

Nagel, L.M. et al. (2017) ‘Adaptive Silviculture for Climate Change: A National Experiment in Manager-Scientist Partnerships to Apply an Adaptation Framework’, Journal of Forestry, 115(3), pp. 167–178. Available at: 10.5849/jof.16-039.

Norberg, A. et al. (2019) ‘A comprehensive evaluation of predictive performance of 33 species distribution models at species and community levels’, Ecological Monographs, 89(3), p. e01370. Available at: 10.1002/ecm.1370.

O’Brien, M.J. et al. (2017) ‘A synthesis of tree functional traits related to drought-induced mortality in forests across climatic zones’, Journal of Applied Ecology, 54(6), pp. 1669–1686. Available at: 10.1111/1365-2664.12874.

Oliva, J., Redondo, M.Á. and Stenlid, J. (2020) ‘Functional Ecology of Forest Disease’, Annual Review of Phytopathology, 58(Volume 58, 2020), pp. 343–361. Available at: 10.1146/annurev-phyto-080417-050028.

Olson, D.M. et al. (2001) ‘Terrestrial Ecoregions of the World: A New Map of Life on Earth: A new global map of terrestrial ecoregions provides an innovative tool for conserving biodiversity’, BioScience, 51(11), pp. 933–938. Available at: 10.1641/0006-3568(2001)051[0933:TEOTWA]2.0.CO;2.

Palik, B.J. et al. (2022) ‘Operationalizing forest-assisted migration in the context of climate change adaptation: Examples from the eastern USA’, Ecosphere, 13(10), p. e4260. Available at: 10.1002/ecs2.4260.

Paulsen, J., Weber, U.M. and Körner, Ch. (2000) ‘Tree Growth near Treeline: Abrupt or Gradual Reduction with Altitude?’, Arctic, Antarctic, and Alpine Research, 32(1), pp. 14–20. Available at: 10.1080/15230430.2000.12003334.

Paz, A., Crowther, T.W. and Maynard, D.S. (2024) ‘Functional and phylogenetic dimensions of tree biodiversity reveal unique geographic patterns’, Global Ecology and Biogeography, 33(9), p. e13877. Available at: 10.1111/geb.13877.

Pecl, G.T. et al. (2017) ‘Biodiversity redistribution under climate change: Impacts on ecosystems and human well-being’, Science, 355(6332), p. eaai9214. Available at: 10.1126/science.aai9214.

Pedlar, J.H. et al. (2012) ‘Placing Forestry in the Assisted Migration Debate’, BioScience, 62(9), pp. 835–842. Available at: 10.1525/bio.2012.62.9.10.

Poggiato, G. et al. (2021) ‘On the Interpretations of Joint Modeling in Community Ecology’, Trends in Ecology & Evolution, 36(5), pp. 391–401. Available at: 10.1016/j.tree.2021.01.002.

Pollock, L.J. et al. (2014) ‘Understanding co-occurrence by modelling species simultaneously with a Joint Species Distribution Model (JSDM)’, Methods in Ecology and Evolution, 5(5), pp. 397–406. Available at: 10.1111/2041-210X.12180.

Pregitzer, K.S. et al. (2000) ‘Responses of tree fine roots to temperature’, New Phytologist, 147(1), pp. 105–115. Available at: 10.1046/j.1469-8137.2000.00689.x.

Quetin, G.R. et al. (2023) ‘Observed forest trait velocities have not kept pace with hydraulic stress from climate change’, Global Change Biology, 29(18), pp. 5415–5428. Available at: 10.1111/gcb.16847.

Rivers, M. et al. (2023) ‘Scientists’ warning to humanity on tree extinctions’, *PLANTS, PEOPLE*, PLANET, 5(4), pp. 466–482. Available at: 10.1002/ppp3.10314.

Roberts, D.R. et al. (2017) ‘Cross-validation strategies for data with temporal, spatial, hierarchical, or phylogenetic structure’, Ecography, 40(8), pp. 913–929. Available at: 10.1111/ecog.02881.

Rosenfeld, J.S. (2002) ‘Functional redundancy in ecology and conservation’, Oikos, 98(1), pp. 156–162. Available at: 10.1034/j.1600-0706.2002.980116.x.

Rueda, M., Godoy, O. and Hawkins, B.A. (2018) ‘Trait syndromes among North American trees are evolutionarily conserved and show adaptive value over broad geographic scales’, Ecography, 41(3), pp. 540–550. Available at: 10.1111/ecog.03008.

Saxe, H. et al. (2001) ‘Tree and forest functioning in response to global warming’, New Phytologist, 149(3), pp. 369–399. Available at: 10.1046/j.1469-8137.2001.00057.x.

Schrodt, F. et al. (2015) ‘BHPMF – a hierarchical Bayesian approach to gap-filling and trait prediction for macroecology and functional biogeography’, Global Ecology and Biogeography, 24(12), pp. 1510–1521. Available at: 10.1111/geb.12335.

Seidl, R. et al. (2017) ‘Forest disturbances under climate change’, Nature Climate Change, 7(6), pp. 395–402. Available at: 10.1038/nclimate3303.

Shapley, L.S. (1953) ‘A value for n-person games’.

Svenning, J.-C. and Sandel, B. (2013) ‘Disequilibrium vegetation dynamics under future climate change’, American Journal of Botany, 100(7), pp. 1266–1286. Available at: 10.3732/ajb.1200469.

Urban, J. et al. (2017) ‘Stomatal conductance increases with rising temperature’, Plant Signaling & Behavior, 12(8), p. e1356534. Available at: 10.1080/15592324.2017.1356534.

Wang, W.J. et al. (2015) ‘Importance of succession, harvest, and climate change in determining future composition in U.S. Central Hardwood Forests’, Ecosphere, 6(12), pp. 1–18. Available at: 10.1890/ES15-00238.1.

Way, D.A. and Oren, R. (2010) ‘Differential responses to changes in growth temperature between trees from different functional groups and biomes: a review and synthesis of data’, Tree Physiology, 30(6), pp. 669–688. Available at: 10.1093/treephys/tpq015.

Wilkes, M.A. et al. (2020) ‘Trait-based ecology at large scales: Assessing functional trait correlations, phylogenetic constraints and spatial variability using open data’, Global Change Biology, 26(12), pp. 7255–7267. Available at: 10.1111/gcb.15344.

Wilkinson, D.P. et al. (2021) ‘Defining and evaluating predictions of joint species distribution models’, Methods in Ecology and Evolution, 12(3), pp. 394–404. Available at: 10.1111/2041-210X.13518.

