## Supplementary Information for "Trait misalignment risk in North American forests under climate change"

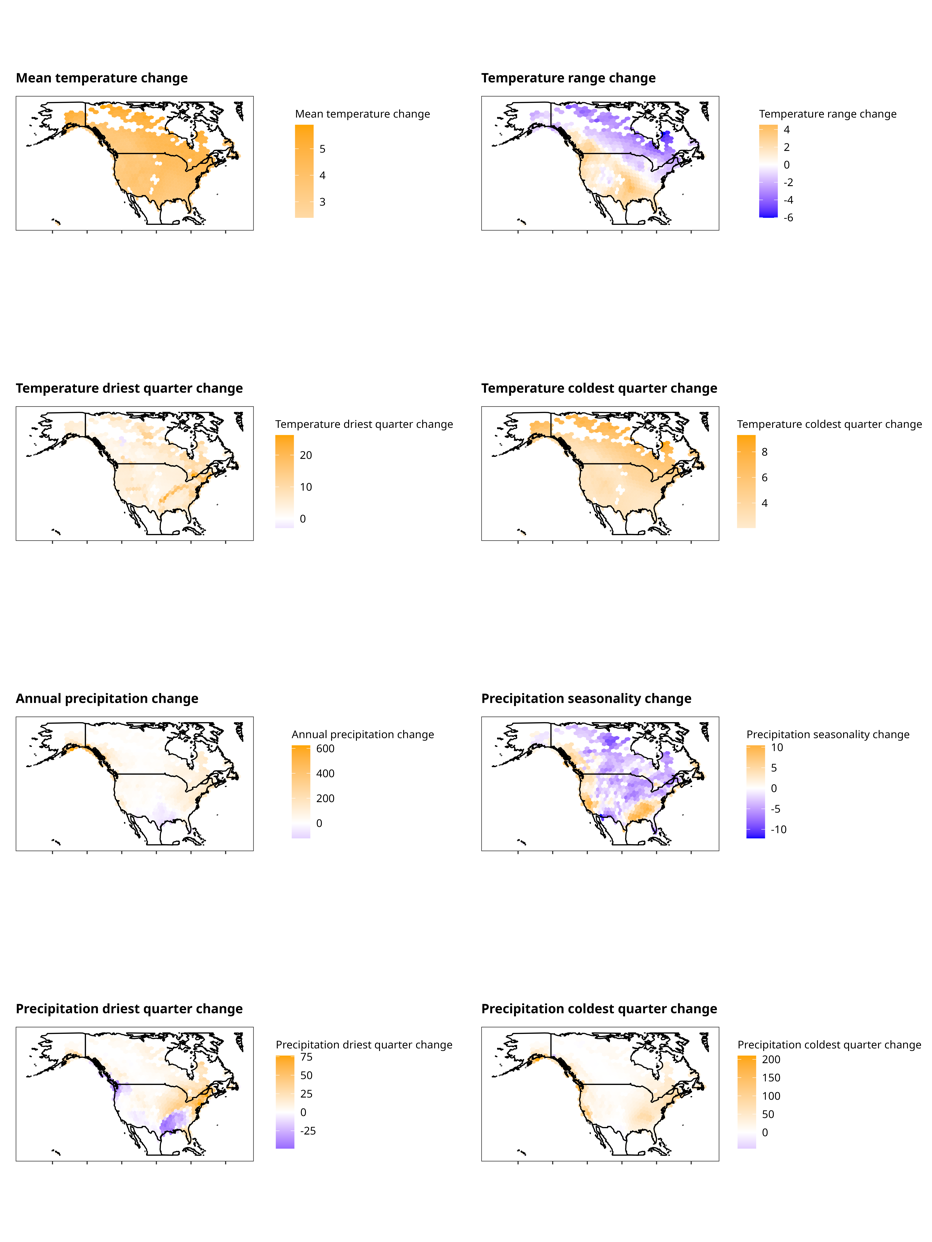

**Supplementary Figure S1: Heatmaps of environmental changes under SSP370.**

*Each hexagonal pixel represents the average absolute change in the environmental variable per plot between current and predicted values under SSP370.*

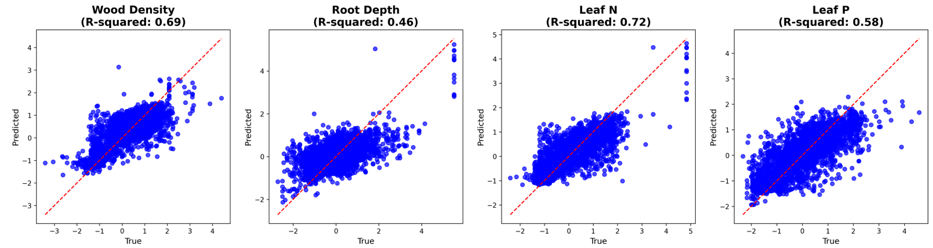

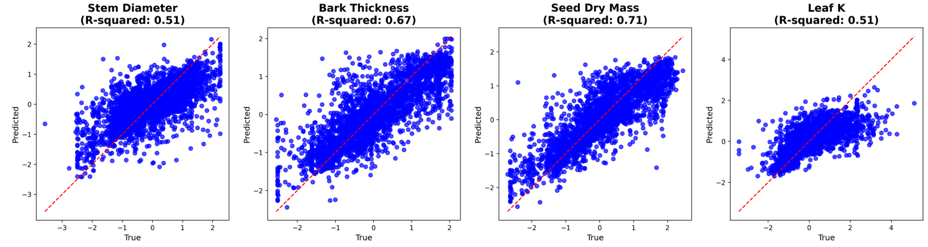

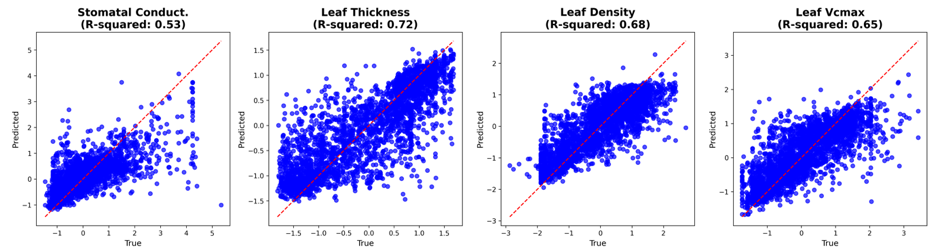

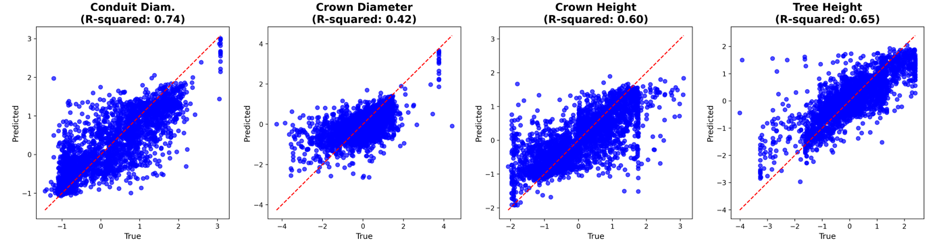

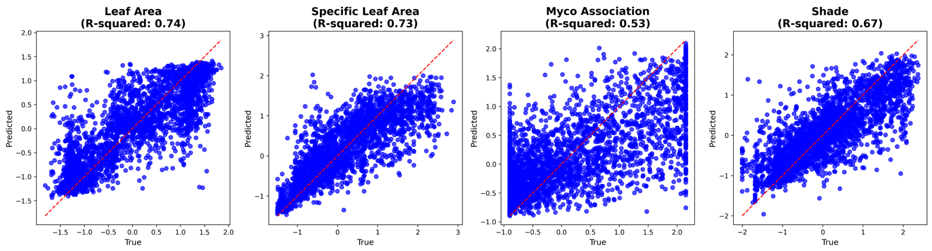

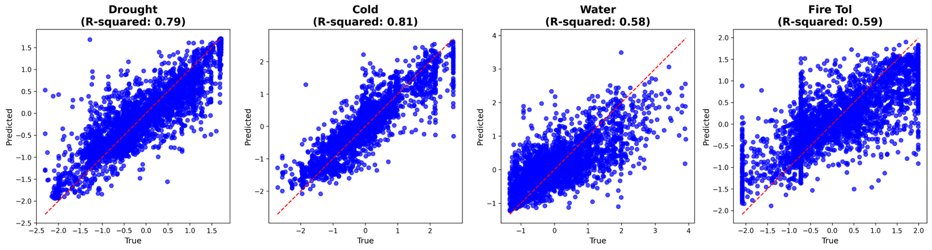

**Supplementary Figure S2: Predicted vs Actual values per trait**

*Multivariate prediction of community plot traits compared with true values. R^2^* *values per trait.*

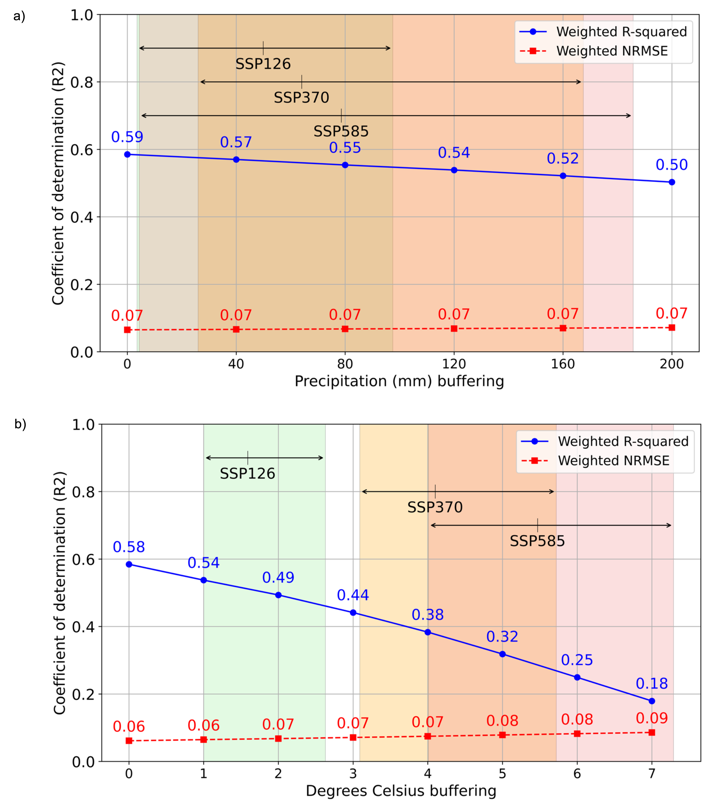

**Supplementary Figure S3: Model forecast horizon**

*(a) Mean annual precipitation and (b) mean annual surface air temperature. The x-axis shows the buffer size for each climate variable, and the y-axis shows model performance (R² and NRMSE). R² values at a buffer size of 0 correspond to standard leave-one-group-out cross-validation performance. Shaded regions indicate the range of projected change in each variable under three SSP scenarios, thereby illustrating how forecast accuracy changes as climates diverge from observed conditions.*

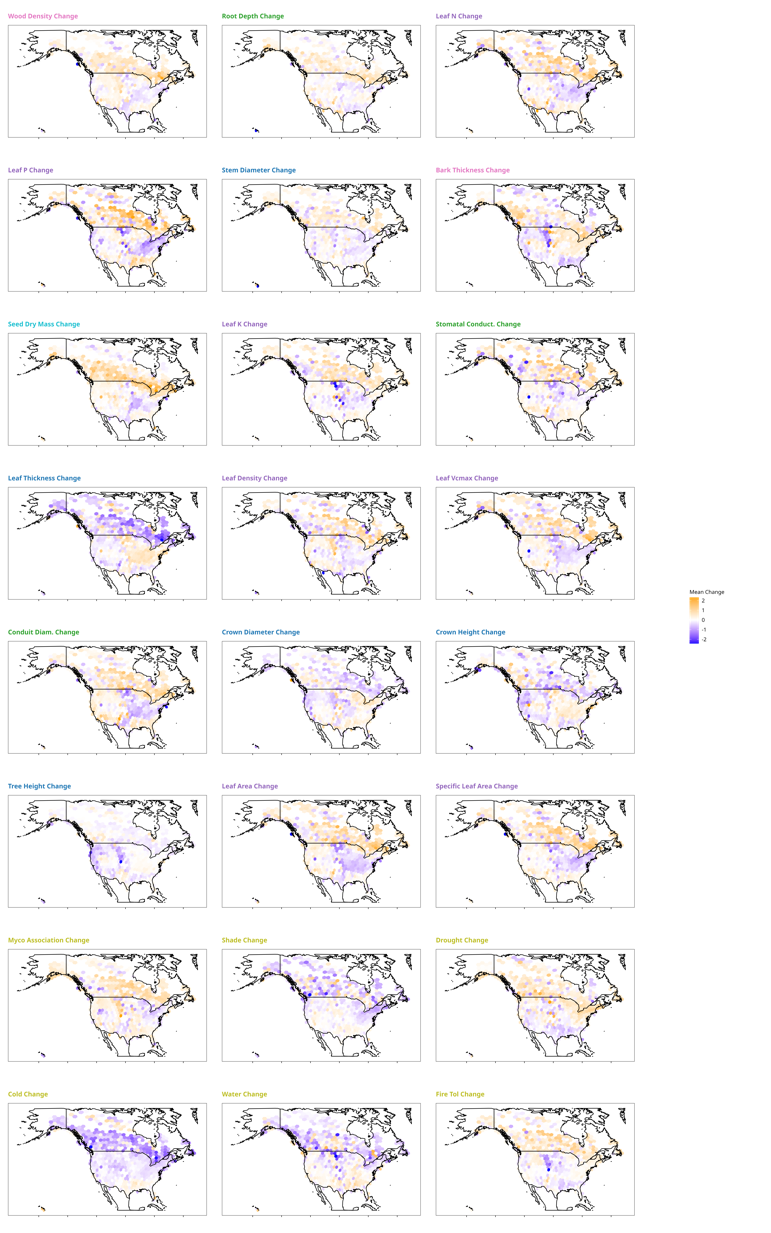

**Supplementary Figure S4: Heatmaps of trait changes under SSP370.** Each hexagonal pixel represents the average absolute change in community-weighted means (CWMs) per plot between current and predicted values under SSP370.

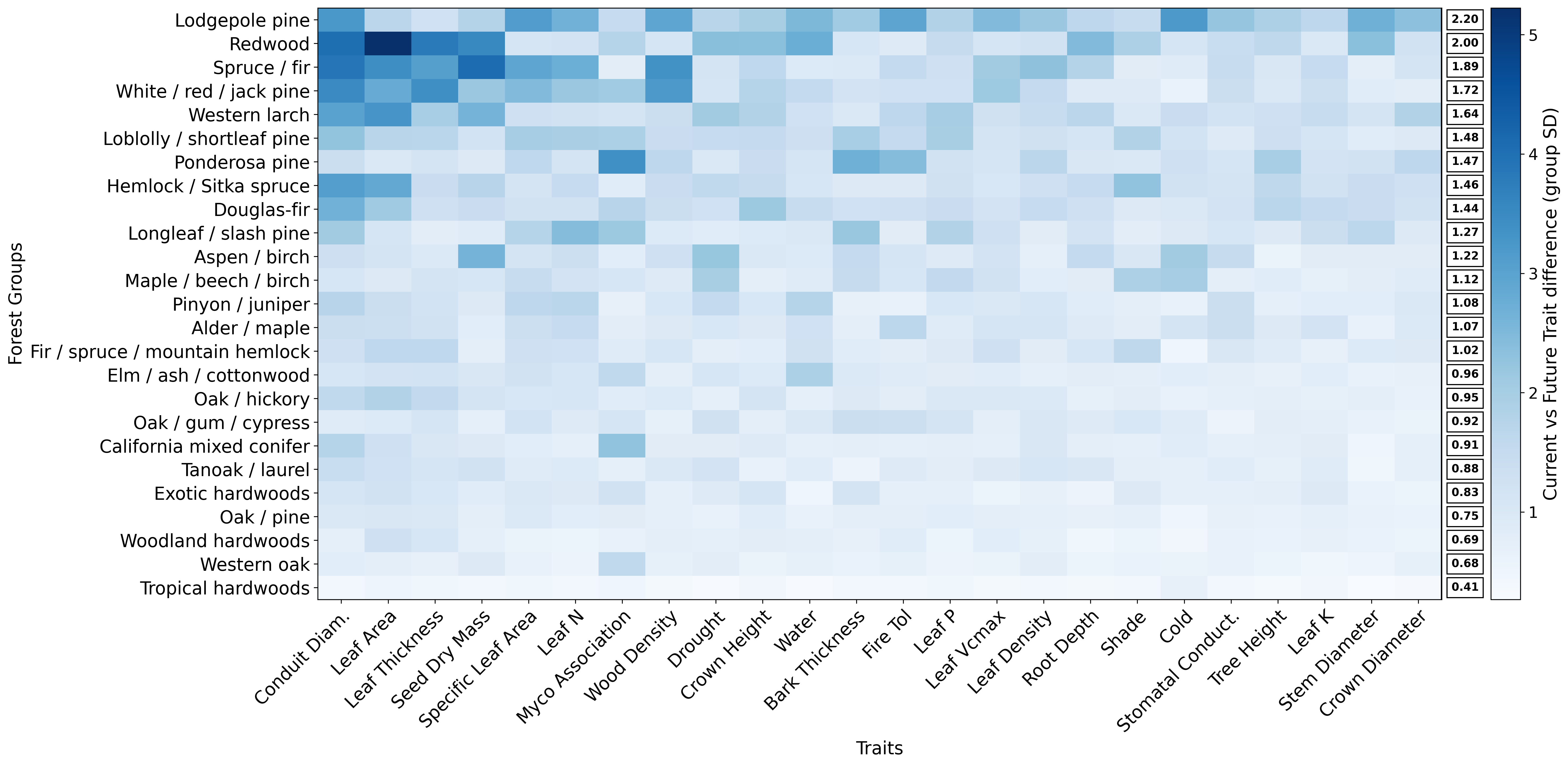

**Supplementary Figure S5: TEM risk by forest group functional composition**

Rows represent forest groups, and columns correspond to functional traits, with colour intensity indicating TEM trait risk for that group — darker shades signify higher risk. Forest groups are sorted by overall TEM risk from highest to lowest, while traits are ordered by their contribution to TEM risk. The final column summarises the average TEM risk per forest group.

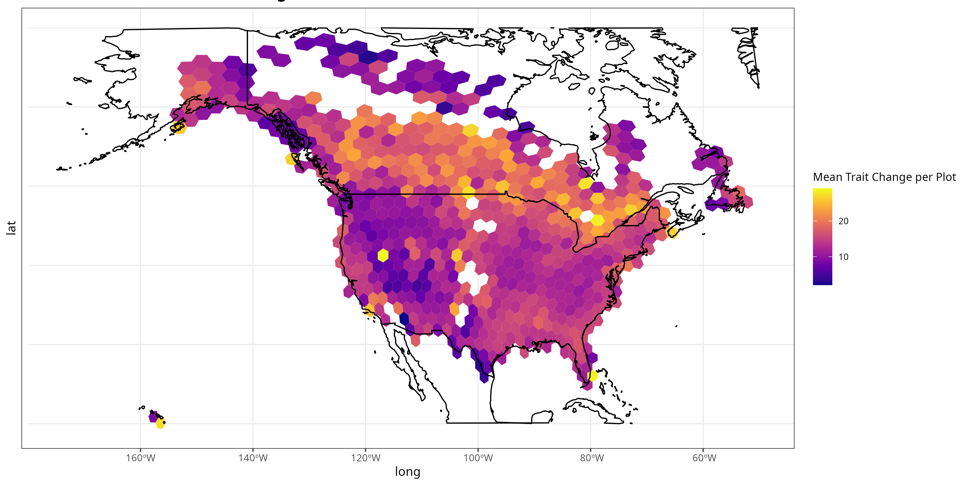

**Supplementary Figure S6: Heatmap of mean trait change under SSP370.** Each hexagonal pixel represents the average absolute change in community-weighted means (CWMs) per plot between current and predicted values under SSP370.

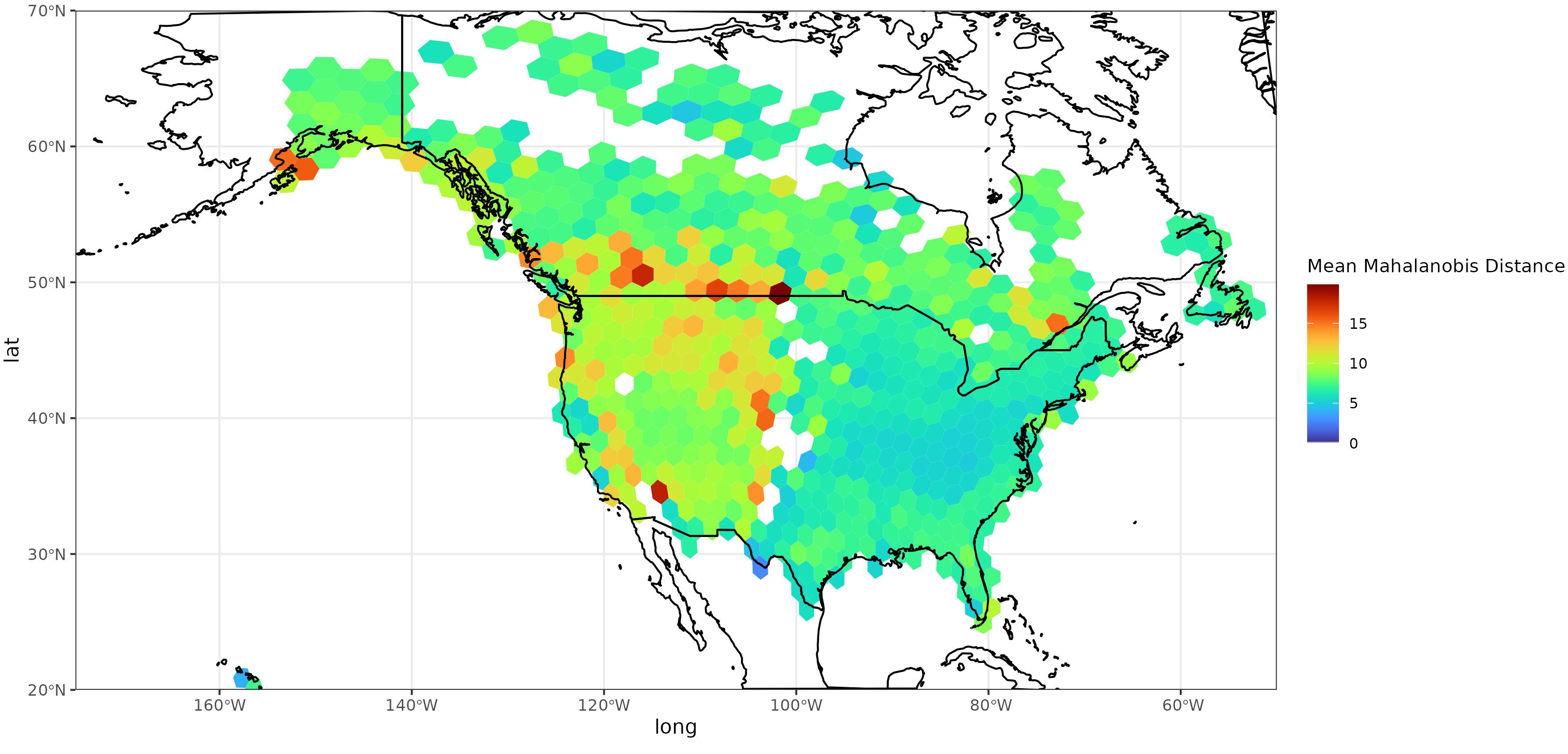

**Supplementary Figure S7: Spatial projections of trait–environment misalignment (TEM) risk.**

*(a) Mean Mahalanobis distance in 24-trait CWM space between present-day and 2100 climate-associated composition (higher values indicate greater overall misalignment) taken from 1000 Monte Carlo draws.*

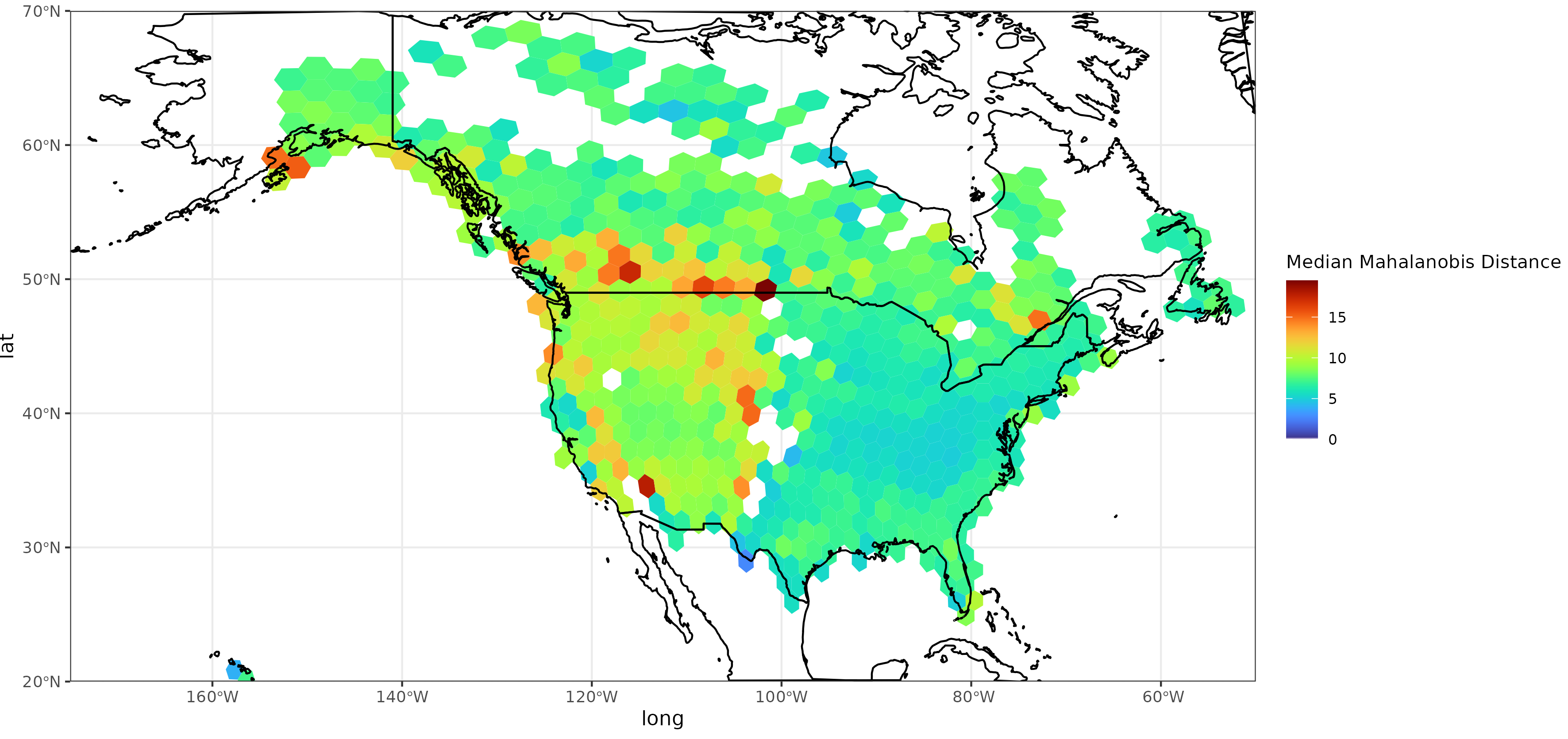

**Supplementary Figure S8: Spatial projections of trait–environment misalignment (TEM) risk.**

*(a) Median Mahalanobis distance in 24-trait CWM space between present-day and 2100 climate-associated composition (higher values indicate greater overall misalignment) taken from 1000 Monte Carlo draws.*

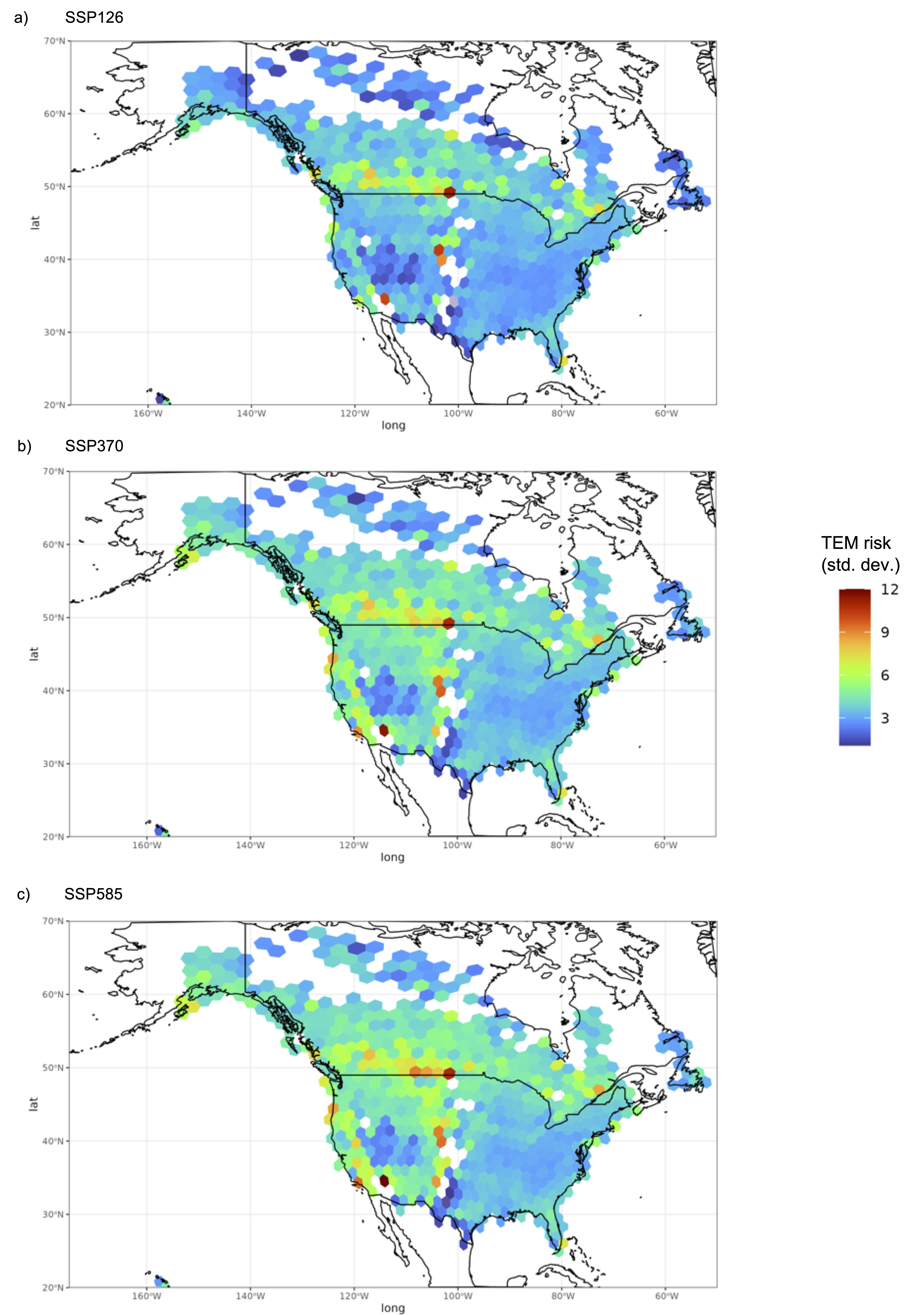

**Supplementary Figure S9: Spatial projections of trait–environment misalignment (TEM) risk across SSPs 126, 370 and 585.**

Each hexbin represents the average number of traits per plot where predicted change exceeds ±3 times the group standard deviation (FV), with a maximum possible value of 24 (total number of traits). a) SSP126; b) SSP370; c) SSP585.

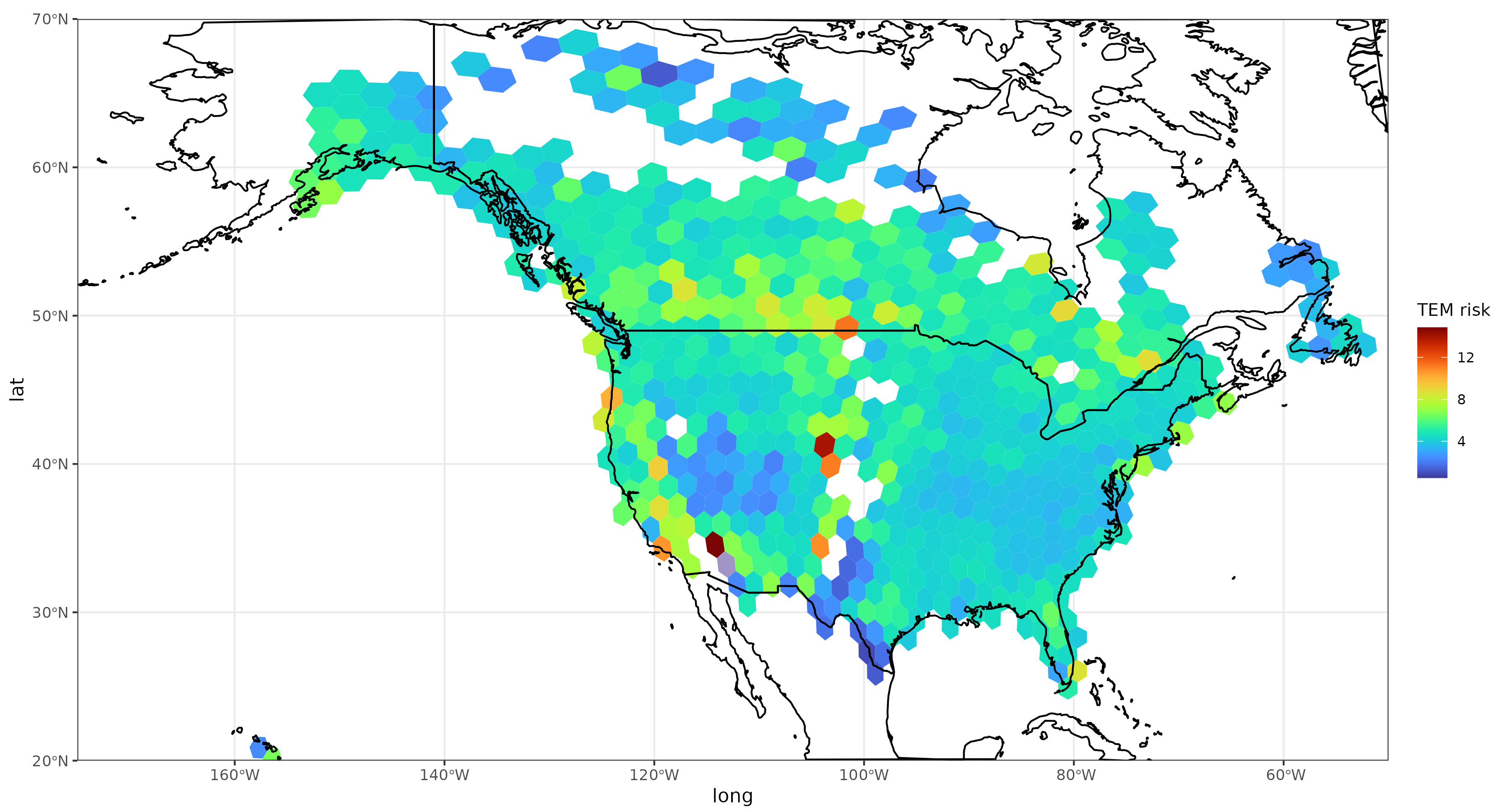

**Supplementary Figure S10: Spatial projections of trait–environment misalignment risk – original CWMs.**

*(a) Mahalanobis distance in 24-trait CWM space between present-day CWMs and 2100 climate-associated composition (higher values indicate greater overall misalignment). No perturbations used – just the original plot CWMs.*

**Supplementary Table S1:** Predicted normalised mean change in trait values across SSPs

| **Trait** | **SSP126** | **SSP370** | **SSP585** |
| --- | --- | --- | --- |
| Bark thickness | 0.018 | 0.001 | 0.021 |
| Cold tolerance | -0.126 | -0.217 | -0.284 |
| Crown diameter, maximum | 0.039 | 0.049 | 0.077 |
| Crown height, maximum | -0.017 | -0.045 | -0.048 |
| Drought tolerance | 0.073 | 0.117 | 0.151 |
| Fire tolerance | 0.04 | 0.032 | 0.051 |
| Leaf area | 0.021 | 0.025 | 0.027 |
| Leaf density | 0.031 | 0.047 | 0.041 |
| Leaf K per mass | 0.04 | 0.064 | 0.058 |
| Leaf N per mass | 0.02 | 0.049 | 0.041 |
| Leaf P per mass | 0.009 | 0.02 | -0.012 |
| Leaf thickness | -0.008 | 0.004 | 0.009 |
| Leaf Vcmax | 0.006 | 0.017 | 0.01 |
| Mycorrhizal association | 0.038 | 0.11 | 0.103 |
| Root depth, maximum | 0.071 | 0.116 | 0.119 |
| Seed dry mass | 0.129 | 0.19 | 0.211 |
| Shade tolerance | -0.082 | -0.147 | -0.183 |
| Specific leaf area | 0.012 | 0.013 | 0.001 |
| Stem conduit diameter | 0.056 | 0.114 | 0.125 |
| Stem diameter | -0.014 | -0.038 | -0.053 |
| Stomatal conductance | 0.029 | 0.109 | 0.114 |
| Tree height, maximum | -0.059 | -0.147 | -0.171 |
| Waterlogging tolerance | -0.007 | 0.03 | 0.022 |
| Wood density | 0.052 | 0.081 | 0.096 |

**Supplementary Table S2:** Wilcoxon Signed-Rank test results on predicted trait changes

|  | **W-statistic** | **p-value** | **Adj. p-value** |
| --- | --- | --- | --- |
| SSP126_vs_Current_wood_density | 398212244 | 1.20E-85 | 8.67E-84 |
| SSP370_vs_Current_wood_density | 376727796 | 1.56E-174 | 1.13E-172 |
| SSP585_vs_Current_wood_density | 368245196 | 2.32E-218 | 1.67E-216 |
| SSP126_vs_Current_root_depth | 399004359 | 5.65E-83 | 4.07E-81 |
| SSP370_vs_Current_root_depth | 378792158 | 1.30E-164 | 9.33E-163 |
| SSP585_vs_Current_root_depth | 377847756 | 4.09E-169 | 2.94E-167 |
| SSP126_vs_Current_leaf_n | 418844069 | 4.20E-30 | 3.02E-28 |
| SSP370_vs_Current_leaf_n | 400919634 | 1.08E-76 | 7.77E-75 |
| SSP585_vs_Current_leaf_n | 411731822 | 5.89E-46 | 4.24E-44 |
| SSP126_vs_Current_leaf_p | 436276909 | 8.19E-06 | 0.000589778 |
| SSP370_vs_Current_leaf_p | 434900632 | 5.50E-07 | 3.96E-05 |
| SSP585_vs_Current_leaf_p | 440937206 | 0.009188118 | 0.661544528 |
| SSP126_vs_Current_stem_diameter | 425497232 | 2.11E-18 | 1.52E-16 |
| SSP370_vs_Current_stem_diameter | 411163512 | 2.26E-47 | 1.63E-45 |
| SSP585_vs_Current_stem_diameter | 404502131 | 1.27E-65 | 9.18E-64 |
| SSP126_vs_Current_bark_thickness | 427182795 | 6.46E-16 | 4.65E-14 |
| SSP370_vs_Current_bark_thickness | 435590596 | 2.21E-06 | 0.000159028 |
| SSP585_vs_Current_bark_thickness | 425634362 | 3.42E-18 | 2.46E-16 |
| SSP126_vs_Current_seed_dry_mass | 350081696 | 0 | 0 |
| SSP370_vs_Current_seed_dry_mass | 318648781 | 0 | 0 |
| SSP585_vs_Current_seed_dry_mass | 310694880 | 0 | 0 |
| SSP126_vs_Current_leaf_k | 374665043 | 9.81E-185 | 7.06E-183 |
| SSP370_vs_Current_leaf_k | 362769795 | 2.80E-249 | 2.02E-247 |
| SSP585_vs_Current_leaf_k | 365044470 | 3.49E-236 | 2.51E-234 |
| SSP126_vs_Current_stomatal_conduct. | 358509531 | 9.23E-275 | 6.65E-273 |
| SSP370_vs_Current_stomatal_conduct. | 300533099 | 0 | 0 |
| SSP585_vs_Current_stomatal_conduct. | 298726010 | 0 | 0 |
| SSP126_vs_Current_leaf_thickness | 438609487 | 0.000413085 | 0.029742087 |
| SSP370_vs_Current_leaf_thickness | 438382752 | 0.000292518 | 0.02106127 |
| SSP585_vs_Current_leaf_thickness | 435090515 | 8.12E-07 | 5.85E-05 |
| SSP126_vs_Current_leaf_density | 419657502 | 1.64E-28 | 1.18E-26 |
| SSP370_vs_Current_leaf_density | 411538546 | 1.95E-46 | 1.41E-44 |
| SSP585_vs_Current_leaf_density | 417035404 | 8.31E-34 | 5.98E-32 |
| SSP126_vs_Current_leaf_vcmax | 433994068 | 7.92E-08 | 5.70E-06 |
| SSP370_vs_Current_leaf_vcmax | 420057269 | 9.59E-28 | 6.90E-26 |
| SSP585_vs_Current_leaf_vcmax | 425328928 | 1.16E-18 | 8.37E-17 |
| SSP126_vs_Current_conduit_diam. | 381036201 | 3.64E-154 | 2.62E-152 |
| SSP370_vs_Current_conduit_diam. | 344746306 | 0 | 0 |
| SSP585_vs_Current_conduit_diam. | 341277302 | 0 | 0 |
| SSP126_vs_Current_crown_diameter | 446196087 | 0.608968302 | 1 |
| SSP370_vs_Current_crown_diameter | 440778123 | 0.00762359 | 0.548898507 |
| SSP585_vs_Current_crown_diameter | 427198084 | 6.79E-16 | 4.89E-14 |
| SSP126_vs_Current_crown_height | 441721411 | 0.021859407 | 1 |
| SSP370_vs_Current_crown_height | 423313909 | 6.56E-22 | 4.72E-20 |
| SSP585_vs_Current_crown_height | 422516887 | 2.86E-23 | 2.06E-21 |
| SSP126_vs_Current_tree_height | 401834667 | 8.81E-74 | 6.34E-72 |
| SSP370_vs_Current_tree_height | 363816617 | 3.30E-243 | 2.37E-241 |
| SSP585_vs_Current_tree_height | 355643266 | 1.31E-292 | 9.46E-291 |
| SSP126_vs_Current_leaf_area | 442177172 | 0.034740085 | 1 |
| SSP370_vs_Current_leaf_area | 437565038 | 7.90E-05 | 0.005689537 |
| SSP585_vs_Current_leaf_area | 436538933 | 1.33E-05 | 0.000954522 |
| SSP126_vs_Current_specific_leaf_area | 406828433 | 6.67E-59 | 4.80E-57 |
| SSP370_vs_Current_specific_leaf_area | 405826244 | 9.44E-62 | 6.80E-60 |
| SSP585_vs_Current_specific_leaf_area | 414698052 | 6.34E-39 | 4.57E-37 |
| SSP126_vs_Current_myco_association | 375359559 | 2.88E-181 | 2.07E-179 |
| SSP370_vs_Current_myco_association | 344147078 | 0 | 0 |
| SSP585_vs_Current_myco_association | 350032570 | 0 | 0 |
| SSP126_vs_Current_shade | 383078751 | 5.91E-145 | 4.26E-143 |
| SSP370_vs_Current_shade | 349772213 | 0 | 0 |
| SSP585_vs_Current_shade | 328374527 | 0 | 0 |
| SSP126_vs_Current_drought | 375834294 | 6.47E-179 | 4.66E-177 |
| SSP370_vs_Current_drought | 345160446 | 0 | 0 |
| SSP585_vs_Current_drought | 322438927 | 0 | 0 |
| SSP126_vs_Current_cold | 331355493 | 0 | 0 |
| SSP370_vs_Current_cold | 268664073 | 0 | 0 |
| SSP585_vs_Current_cold | 222929897 | 0 | 0 |
| SSP126_vs_Current_water | 419942496 | 5.79E-28 | 4.17E-26 |
| SSP370_vs_Current_water | 397067136 | 1.39E-89 | 1.00E-87 |
| SSP585_vs_Current_water | 400656178 | 1.53E-77 | 1.10E-75 |
| SSP126_vs_Current_fire_tol | 401091003 | 3.82E-76 | 2.75E-74 |
| SSP370_vs_Current_fire_tol | 399653946 | 8.14E-81 | 5.86E-79 |
| SSP585_vs_Current_fire_tol | 389029855 | 9.36E-120 | 6.74E-118 |
